# Mapping and engineering RNA-controlled architecture of the multiphase nucleolus

**DOI:** 10.1101/2024.09.28.615444

**Authors:** SA Quinodoz, L Jiang, AA Abu-Alfa, TJ Comi, H Zhao, Q Yu, LW Wiesner, JF Botello, A Donlic, E Soehalim, C Zorbas, L Wacheul, A Košmrlj, DLJ Lafontaine, S Klinge, CP Brangwynne

## Abstract

Biomolecular condensates are key features of intracellular compartmentalization. As the most prominent nuclear condensate in eukaryotes, the nucleolus is a layered multiphase liquid-like structure and the site of ribosome biogenesis. In the nucleolus, ribosomal RNAs (rRNAs) are transcribed and processed, undergoing multiple maturation steps that ultimately result in formation of the ribosomal small subunit (SSU) and large subunit (LSU). However, how rRNA processing is coupled to the layered nucleolar organization is poorly understood due to a lack of tools to precisely monitor and perturb nucleolar rRNA processing dynamics. Here, we developed two complementary approaches to spatiotemporally map rRNA processing and engineer *de novo* nucleoli. Using sequencing in parallel with imaging, we found that rRNA processing steps are spatially segregated, with sequential maturation of rRNA required for its outward movement through nucleolar phases. Furthermore, by generating synthetic *de novo* nucleoli through an engineered rDNA plasmid system in cells, we show that defects in SSU processing can alter the ordering of nucleolar phases, resulting in inside-out nucleoli and preventing rRNA outflux, while LSU precursors are necessary to build the outermost layer of the nucleolus. These findings demonstrate how rRNA is both a scaffold and substrate for the nucleolus, with rRNA acting as a programmable blueprint for the multiphase architecture that facilitates assembly of an essential molecular machine.

## Introduction

Biomolecular condensates have emerged as a ubiquitous feature of cellular compartmentalization^1–3^. These include various nuclear bodies, like nucleoli and nuclear speckles, which concentrate DNA, RNA, and proteins of shared functions such as transcription, and RNA processing. There are also cytoplasmic structures, such as signaling clusters, P bodies, and stress granules, associated with various other functions. Condensates form through phase separation and related phase transitions, which result from a competition between entropy and molecular interaction energies of multivalent biomolecules^4,5^. Among their protein and nucleic acid components, RNA plays a particularly central role in promoting and modulating condensate assembly^6^. RNA-related functions may also be altered by the physicochemical environment within condensates^7,8^, but dissecting this relationship between condensate structure and the biochemical reactions occurring within them has been a key challenge for the field.

The nucleolus provides an ideal model for studying condensate structure-function relationships, particularly in the context of RNA processing. The nucleolus is a multiphase organelle composed of three nested subcompartments - the innermost fibrillar center (FC), middle dense fibrillar component (DFC), and outer granular component (GC)^9–11^ (**Figure 1A**). Various biochemical reactions, including the transcription, folding, cleavage and modifications of precursor ribosomal RNA (pre-rRNA) and assembly of pre-ribosomal ribonucleoproteins (RNPs), are initiated within this fascinating multiphase structure. Pre-rRNA is transcribed by RNA Pol I at the FC/DFC boundary and moves radially outwards as intermediates of small and large ribosomal subunits (SSU and LSU) are formed^12–14^. As part of this process, small nucleolar RNA (snoRNA)-guided snoRNPs and other enzymes place hundreds of RNA modifications (e.g., 2’-O-methylation and pseudouridylation) at specific sites, fold pre-rRNA, and sequentially cleave the ∼13 kb pre-rRNA transcript into three mature ribosomal RNAs, 18S (1.9 kb), 5.8S (0.15 kb), and 28S rRNAs (5 kb), to assemble the SSU and LSU of the ribosome^15–17^. These transcription and processing steps are intimately coupled to nucleolar structure: RNA Pol I inhibition causes a major rearrangement of the nucleolar phases known as “nucleolar caps”^18–20^, while drug treatments perturbing rRNA processing or the knockdown of ribosome biogenesis factors result in altered nucleolar morphology^21–23^.

**Figure 1:**
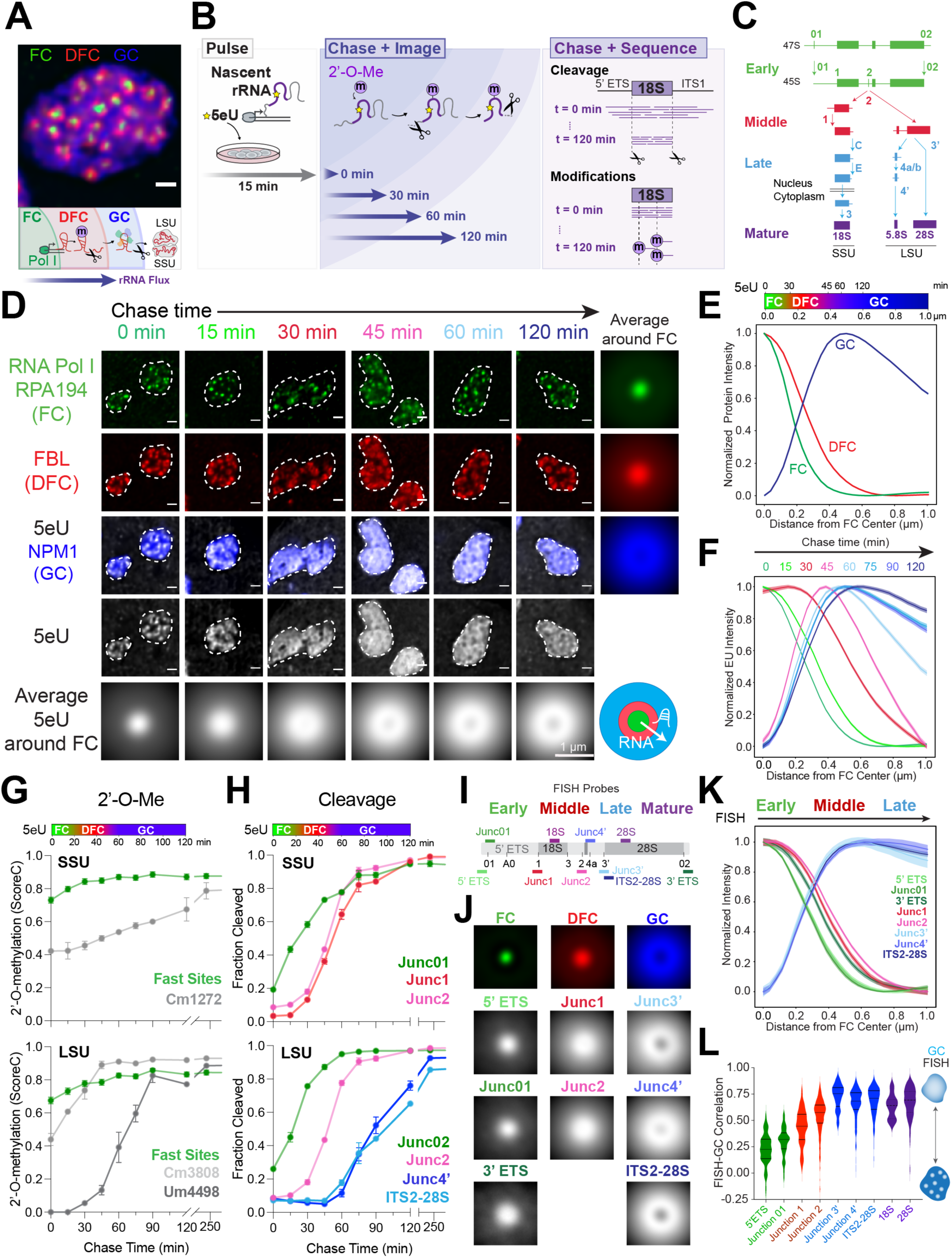
Sequencing and imaging of nascent ribosomal RNA flux provides a spatiotemporal map of processing in the nucleolus. **(A)** The three phases of the nucleolus in MCF10A cells: the fibrillar center (FC, green), dense fibrillar component (DFC, red), and granular component (GC, blue). Schematic of pre-rRNA processing and outflux as it is assembled into small and large ribosomal subunits (SSU and LSU). **(B)** The 5eU-seq and 5eU-imaging approach. Cells were pulsed with 5eU to label nascent RNA for 15 min, followed by chase with excess uridine over various time points for imaging to measure rRNA flux or sequencing to measure rRNA cleavage and 2’-*O*-methylation (2’-*O*-Me). **(C)** Schematic of cleavage steps occurring during pre-rRNA processing (only one of the two cleavage pathways is drawn for simplicity). Cleavage junctions and intermediates are categorized into early (green), middle (red), late (blue), and mature (purple) based on 5eU-seq data in H. **(D)** 5eU-imaging in MCF10A cells shows radial outflux of 5eU-labeled pre-rRNA from the FC center over different chase times. Individual nucleoli are outlined by dashed lines. Averaged 5eU images around FCs are shown. **(E)** Min-max normalized intensities (y axis) of FC, DFC, GC nucleolar phases by distance from FC center (μm, x axis), quantified from images in D; number of nucleoli (n) = 4274. A color bar indicating the localization of FC, DFC, GC phases is shown. **(F)** Min-max normalized intensities of 5eU (y axis) by distance from FC center (μm, x axis) over different chase timepoints, quantified from images in d; number of nucleoli (n) = 717, 459, 499, 603, 470, 451, 550, 525 for sequential time points. **(G)** 18S and 28S pre-rRNA 2’-*O*-methylation (2’-*O*-Me) levels (ScoreC) detected by 5eU-seq over time. The top color bar relates chase time with distance, generated based on measured 5eU peak location over time (Supplementary Figure 2A). **(H)** Fraction of pre-rRNA cleaved at early, middle, and late sites over time measured by 5eU-seq. **(I)** Schematic of RNA FISH probe localization along rDNA. **(J)** Average fluorescence images around FCs of all FISH probes and the three nucleolar phases. See Supplementary Figure 2D for example images. **(K)** Min-max normalized intensity of FISH probes over distance from FC center with probes for early, middle, late rRNA cleavage junctions. **(L)** Quantified Pearson’s correlation between all FISH probes and GC. For k-l, the number of nucleoli (n) = 72 (5’ ETS), 95 (3’ ETS), 24 (Junc01), 111 (Junc1), 230 (Junc2), 38 (Junc3’), 105 (Junc4’), ITS2-28S (318), 18S (72), 28S (310). Violin plots are centered by median and quartiles are shown. All scale bars = 1 μm. All error bars are s.e.m. All FCs are labeled with RNA polymerase I (Pol I) subunit RPA194 IF, DFCs with fibrillarin (FBL) IF and GC with endogenously tagged mTagBFP2-NPM1 as validated in Supplementary Figure 2E-F.

The coupling between pre-rRNA processing at the molecular scale and the emergent multiphase nucleolar morphology at the micrometer scale has thus far remained elusive. “Bottom-up” *in vitro* reconstitution approaches utilize purified nucleolar proteins and rRNAs to form nucleoli mimics^11,24,25^, but they are vast simplifications of the complexity of real nucleoli. Within cells, pre-rRNA processing has been measured using pulse-chase radiolabeling or Northern blot analysis since the discovery of ribosome assembly in human cells^26,27^, but these approaches lack spatial information of where these events occur in the nucleolus. Instead, RNA fluorescence in situ hybridization (FISH) has been employed for spatial visualization^28,29^, but it cannot measure rRNA modifications or resolve the kinetics of rRNA processing events. Thus, it has been challenging to directly link temporal dynamics of rRNA processing with its spatial movement through the nucleolus. Additionally, the repetitive nature of rDNA genes has limited the ability to mutate it endogenously to directly study how rRNA contributes to nucleolar morphology^30–32^. Progress in this field is thus currently limited by the absence of techniques for both mapping where and when these specific pre-rRNA processing steps occur and dissecting how they contribute to the formation and spatial organization of each phase of the nucleolus in living cells.

Here, we develop two complementary approaches that address these challenges and reveal new insights into the relationship between rRNA processing and nucleolar structure. First, we introduce a method performing sequencing in parallel with imaging to precisely measure the kinetics of pre-rRNA cleavage and modification with single nucleotide resolution and map where they take place inside the nucleolus. We find that rRNA processing steps occur in a spatially segregated manner within the nucleolus, with sequential maturation of rRNA required for its outward movement through nucleolar phases. Most small subunit assembly steps occur within the DFC phase, while large subunit assembly steps occur in both the DFC and GC phases. To molecularly dissect the contributions of rRNA sequence to nucleolar structure, we deploy an engineered rDNA plasmid system to construct a synthetic nucleolus in cells. By mutating rDNA, we find that LSU precursors are necessary for the formation of the GC phase. Surprisingly, perturbing SSU pre-rRNA processing in both synthetic and endogenous nucleoli leads to a remarkable “inversion” of nucleolar phases, establishing an essential role of SSU processing in maintenance of normal multiphase organization. Finally, to understand how rRNA processing regulates nucleolar morphology, we develop a mathematical framework and modeling approach which provides a simple physical mechanism of how altered concentrations of rRNA intermediates could change interfacial tensions to shape the multiphase nucleolus. Taken together, our study provides a detailed dissection of how different rRNA species build and arrange the nucleolar phases and reveal how the multiphase architecture contributes to proper rRNA flux and ribosome assembly.

## Results

### Sequencing and imaging of nascent ribosomal RNA flux provides a spatiotemporal map of processing in the nucleolus

To measure rRNA processing in space and time, we developed a 5-ethynyl uridine (5eU) nucleotide analog-based pulse-chase labeling approach to measure both nucleolar RNA localization, by super resolution fluorescence microscopy, and processing state, by RNA-sequencing (**Figure 1B**). This approach requires modifying existing nascent RNA-seq strategies^33^ to remove background arising from the high abundance of mature rRNA in ribosomes, which we found can impair sensitive detection of nascent pre-rRNA processing (**Supplementary** Figure 1A-D). Briefly, we labeled nascently-transcribed RNA for 15 minutes (pulse) with the 5eU nucleotide analog and then chased this nascent RNA population with excess unlabelled uridine over time. Because 5eU nucleotides can be conjugated with a dye using click chemistry (5eU-imaging), we can visualize the radial outflux of nascently transcribed rRNA from the FC/DFC interface, where RNA Pol I is synthesizing rRNA, to the outer GC layer over time (**Figure 1B**)^34^. Alternatively, 5eU nucleotides can be conjugated with biotin, enabling purification of these nascently transcribed RNAs with streptavidin beads followed by deep sequencing (5eU-seq)^33^ to measure both rRNA cleavage and modification (2’-*O*-methylation) steps in one experiment (**Supplementary** Figure 1A-D, **H-I)**. Importantly, 5eU incorporation does not impair pre-rRNA synthesis, pre-rRNA processing, or modification significantly (**Supplementary** Figure 1E-G).

Using this approach, we can quantitatively measure both rRNA localization (**Figure 1D-F**) and rRNA processing (**Figure 1C, G-H**) over time. We find that rRNA cleavage steps correlate with movement of rRNA through the nucleolar phases (**Figure 1D-F, H and** **Supplementary** Figure 2A-B). Specifically, early cleavage steps (01 and 02 junctions) begin immediately after transcription (∼0-15 min) (**Figure 1H**), which coincides with 5eU rRNA localization near the FC/DFC boundary at early chase time points (**Figure 1D-F**), consistent with these cleavage steps occurring co-transcriptionally or immediately after transcription^16,35^. Next, we observe subsequent cleavage steps (1 and 2 junctions) as rRNA fluxes from the DFC to GC (∼30-45 min), and late cleavages occur as RNA fluxes from the GC to nucleoplasm (∼60-90 min) (**Figure 1D-F, H**). This spatiotemporal map suggests that nucleolar SSU (18S rRNA) processing steps may be largely completed before RNA enters the GC, while LSU (5.8S/28S rRNA) processing steps may occur throughout the nucleolus as they take markedly longer (**Figure 1H**). This is consistent with pioneering early work in yeast showing that nuclear dwelling time of SSU precursors is far less than those of LSU^36^. In addition, we find that the vast majority of 2’-*O*-methylations occur within 15 minutes after transcription (**Figure 1G**), consistent with findings that snoRNA-mediated 2’-*O-* methylations are largely placed co-transcriptionally in yeast^35^. However, a handful of post-transcriptional 2’-*O-*methylations (e.g., 28S Um4498, Gm4499) occur significantly later (∼60-90min), corresponding to when 5eU rRNA moves into the GC phase and consistent with LSU processing taking longer in the nucleolus (**Figure 1G and** **Supplementary** Figure 1H-I).

To validate the 5eU-seq and imaging approach, we performed RNA FISH (**Figure 1I-L and** **Supplementary** Figure 2C-D) using probes spanning several cleavage junctions. The rationale was that, since probes can no longer hybridize after cleavage, the distance between the FC and where uncleaved species are no longer detected indicates where processing occurred. RNA FISH shows early pre-rRNA species (01 junction) at the FC/DFC boundary, consistent with prior work^28^. Next, junctions 1 and 2 are primarily localized within the DFC and at the DFC/GC boundary, consistent with their cleavages occurring at the same time scale as 5eU rRNA moves from the DFC into the GC (∼30min) (**Figure 1H**). Late LSU cleavage steps (e.g. ITS2), which occur (∼60min) after 5eU RNA enters the GC, are indeed localized within the GC (**Figure 1J-L**), further demonstrating that some LSU processing steps occur in the GC, while most SSU processing steps are completed before RNA enters the GC (**Figure 1H**). Using a kinetic model to deconvolve the contributions of individual intermediates, these FISH data could be directly compared to our 5eU-seq and imaging data, with the resulting spatial distributions comparable between the two (**Supplementary Note 1**). Taken together, this map of where different rRNA processing steps occur demonstrates that cleavage steps are spatially segregated in different phases of the nucleolus, with sequential RNA processing coinciding with outward movement of RNA through nucleolus.

### Impaired rRNA processing impacts rRNA flux and alters nucleolar morphology

Our findings above show that rRNA processing is correlated with its spatial outflux; however, it is unclear if these processing steps are strictly required for rRNA movement through the nucleolus. To test this, we initially perturbed rRNA processing with chemical inhibitors or knockdowns of processing factors and measured the impact on rRNA processing, and flux, as well as examining potential changes to nucleolar morphology (**Figure 2A**). First, we treated cells with a CDK9 inhibitor, flavopiridol (FVP), which blocks rRNA processing but not transcription^22^. Using 5eU-seq to label nascently transcribed rRNA after cells were treated with FVP or DMSO for 1 hour, we find that FVP treatment results in major impairment of all but one rRNA cleavage step - only the earliest co-transcriptional 01 junction proceeds normally (**Figure 2B and** **Supplementary** Figure 3A-C). Surprisingly, while 5eU-seq reveals that all 2’-O-Me levels are significantly reduced by FVP, we find that SSU and LSU are differentially impacted, with modifications on 28S reduced more than those on 18S (**Figure 2D and** **Supplementary** Figure 4A-D). Consistent with prior studies, FVP-treated nucleoli form a typical “necklace” morphology visible in confocal fluorescence microscopy^22^, where the outer GC phase appears “detached” from the inner FC/DFC phases (**Figure 2E and** **Supplementary** Figure 5C).

**Figure 2:**
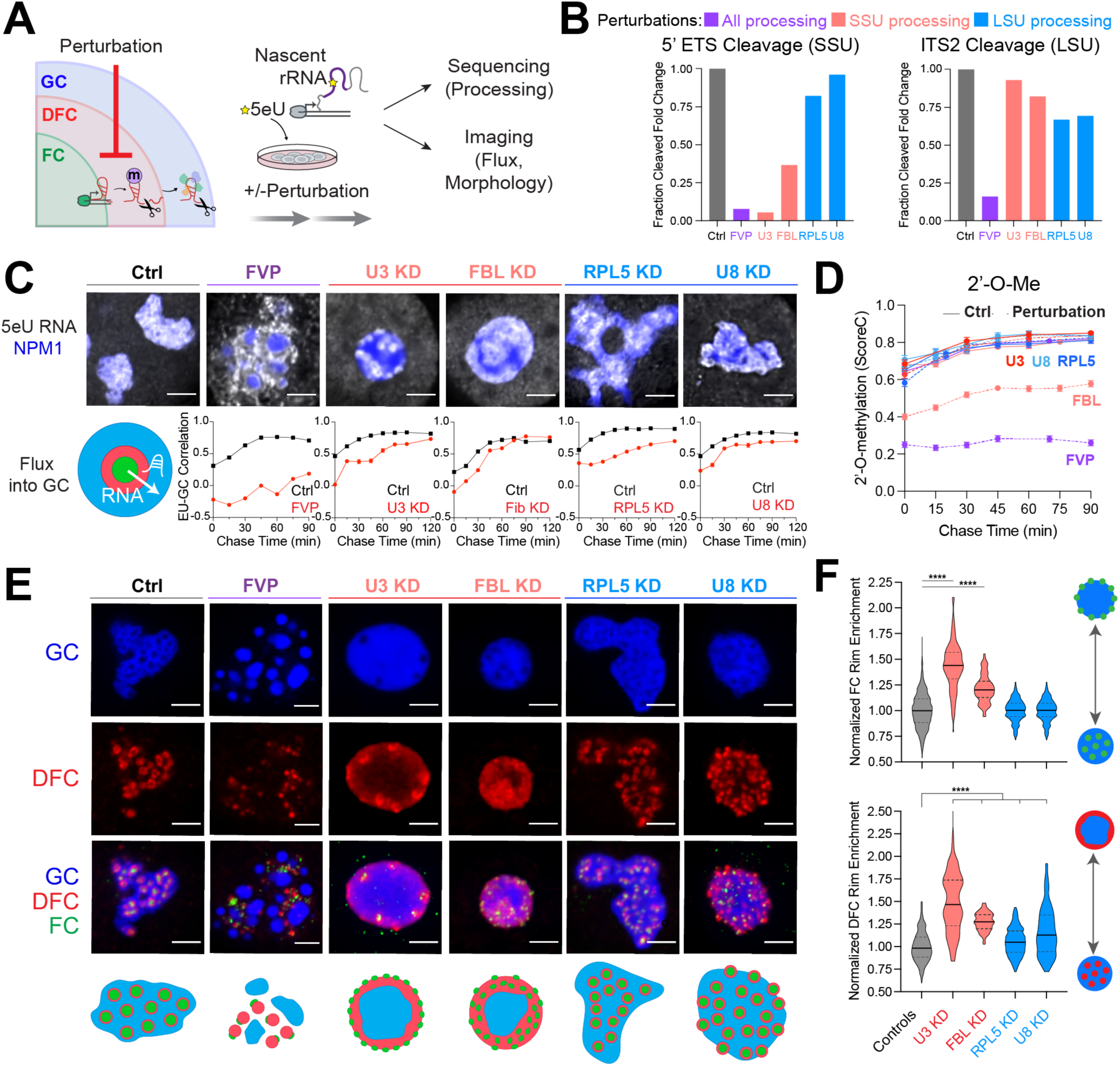
Impaired rRNA processing impacts rRNA flux and alters nucleolar morphology. **(A)** Cells were treated with various pre-rRNA processing perturbations followed by 5eU pulse chase. 5eU-seq measures defects in pre-rRNA processing while 5eU imaging measures the flux of nascent RNA through the nucleolus as well as nucleolar morphology changes. **(B)** Fold change in 5’ ETS cleavage (junction 1) or ITS2 (region downstream of 3’ junction) measured using 5eU-seq upon different perturbations: 2 μM Flavopiridol (FVP) treatment (general processing inhibitor, purple), knockdown (KD) of SSU processing factors (U3 snoRNA or FBL, red), or LSU processing factors (RPL5 or U8 snoRNA, blue). **(C)** Representative images of 5eU-labeled RNA (white) in nucleolus (mTagBFP2-NPM1) after 60 min chase in control and in various perturbation conditions. Quantification of 5eU correlation with NPM1 (GC) 0-120 minutes after transcription (chase time) upon all perturbations. **(D)** Average 18S and 28S rRNA 2’-*O*-Me levels (quantified by ScoreC) by 5eU-seq upon all perturbations (dashed lines) or control (solid lines) across 0-90 min after transcription (chase time). **(E)** Representative images and schematics of GC (NPM1, blue), DFC (FBL, red), and FC (RPA194, green) labeled with IF for all conditions except for FBL KD, where mTagBFP2-NPM1, NOP56-mCherry, RPA16-GFP was used. **(F)** Quantification of FC and DFC enrichment on the rim of the nucleolus in control and all perturbations (normalized to control average value). Violin plots are centered by median (solid line) and quartiles are shown (dashed lines). **** p-value < 0.0001 (two-tailed Mann-Whitney test) n = 1116 (Controls), 97 (Fib), 228 (RPL5), 153 (U3), 105 (U8) cells. All scale bars = 3 μm. All error bars are s.e.m.

Given that FVP results in inhibition of almost all processing steps, we pretreated cells with FVP for 1 hour and then performed pulse-chase 5eU-imaging to determine whether these unprocessed rRNA species can flux into the GC phase. While rRNAs normally move into the GC after a 60-minute chase in the DMSO-control cells (**Figure 2C**), unprocessed rRNAs in the FVP-treated cells accumulate around the GC surface, coming into close contact with the boundary but not partitioning into the GC (**Figure 2C and** **Supplementary** Figure 5E); RNA FISH again validates these findings (**Supplementary** Figure 5A-B). Supporting this, FVP washout experiments demonstrated that restarting pre-rRNA processing led to the reattachment of the GC phase (labeled by NPM1) with FCs and DFCs within ∼30-60 mins (**Supplementary** Figure 5D **and Supplementary Video 1-2**). This reattachment is consistent with the timing of when nascently transcribed rRNA is normally cleaved and enters the GC (**Figure 1D**); this suggests that as pre-rRNA processing resumes, processed rRNAs can now partition into the outer GC layer. Taken together, our results suggest that rRNA processing is required for the selective flux of processed rRNAs into the GC, and that this processing-gated movement of RNAs plays a key role in shaping nucleolar morphology.

Since FVP is a blunt perturbation affecting most rRNA processing steps, we next examined the consequences on rRNA flux upon more precise knockdowns of representative rRNA processing factors involved in specific rRNA processing steps. We started by using antisense oligonucleotides (ASOs) to deplete the box C/D snoRNA U3^37^, a component of the SSU processome containing about 50 nucleolar SSU assembly factors involved in SSU processing, including 5’ ETS cleavage^37,38^. In control cells, our 5eU-seq approach clearly reveals the sequential processing of 5’ ETS over the course of 90 min. Upon U3 depletion, we indeed observe disruptions of the two 5’ ETS (01 and 1) cleavage junctions in the SSU processing pathway (**Figure 2B and** **Supplementary** Figure 3D), as expected^37^. Interestingly, we observe that the rate of junction 1 cleavage is almost fully stopped, while the co-transcriptional 01 junction still proceeds, but is much delayed (**Supplementary** Figure 3D). This is consistent with expectations because junction 1 cleavage (by Utp24 within the SSU processome) cannot occur in the absence of U3 base pairing with the 5’ ETS, while 01 cleavage is mediated by XRN2 and MTR4^38–40^. Conversely, rates of LSU processing are not impacted upon U3 ASO treatment, consistent with previous work^37^ (**Figure 2B and** **Supplementary** Figure 3D). Given that rRNA processing rates are reduced, we then performed 5eU-imaging and found that the rate of RNA flux into the GC is similarly slowed (**Figure 2C**). Altogether, these data show that 5eU-seq can be used to quantitatively measure rRNA cleavage and modification rates at single nucleotide resolution and, when coupled with 5eU imaging, reveal that pre-rRNA processing rates are tightly connected to the rates of rRNA movement through the nucleolus.

### Perturbed SSU processing results in an “inside-out” nucleolar morphology

We next examined the consequences of these processing defects upon U3 knockdown on the overall morphology of the nucleolus. Remarkably, U3 knockdown results in an “inside-out” morphology where the ordering of the 3 nucleolar phases are reversed, and the FC and DFC phases move towards the periphery of the nucleolus (**Figure 2E-F and** **Supplementary** Figure 6A-F). This is distinct from the well-known “nucleolar cap” morphology^19,22^, where FCs also move toward the periphery of the nucleolus, but fuse due to transcriptional inhibition (**Supplementary** Figure 6B). Instead, the number of FCs remains high upon U3 snoRNA knockdown because of active Pol I transcription (**Supplementary** Figure 6B). We also observe the number of nucleoli reduces to one upon U3 snoRNA knockdown (**Supplementary** Figure 6G).

We next examined if the nucleolar inversion could be phenocopied by knocking down another component of the U3 snoRNP, fibrillarin (FBL) (**Supplementary** Figure 6H). 5eU-seq reveals that fibrillarin knockdown slows SSU cleavage and broadly reduces modifications (**Figure 2B,D and** **Supplementary** Figure 3E, **4D-E**), consistent with its function as a 2’-*O*-methyltransferase and component of the SSU processome^38,41^. Interestingly, rates and levels of 28S modifications change more significantly than 18S rRNA modifications, except for the 4499-Gm modification on 28S rRNA, not placed by fibrillarin^42^, which is unaffected (**Supplementary** Figure 4D-E). We find that fibrillarin KD also results in a partially inverted morphology, like that seen with U3 KD (**Figure 2E-F**), underscoring the importance of proper SSU processing in nucleolar morphology. Inversion is not a generic trait of processing defects, as depletion of RPL5 (uL18) or the box C/D snoRNA U8 result in distinct nucleolar morphologies (**Figure 2E-F and** **Supplementary** Figure 6I-K). We note that these RPL5 and U8 knockdowns primarily impact LSU processing, with some additional effects in the SSU pathway, consistent with prior studies^23,37^ (**Figure 2B and** **Supplementary** Figure 3F-G). In addition to the slowed pre-rRNA processing rates, the rate of rRNA flux into the GC phase is similarly reduced for all perturbations (**Figure 2C and** **Supplementary** Figure 7). Altogether, these findings suggest that rRNA processing both shapes nucleolar morphology and impacts the rate of rRNA flux into the GC.

### Engineered synthetic nucleoli in cells recapitulate normal multiphase architecture

To gain further insight into how pre-rRNA processing shapes nucleolar morphology, we sought to generate an engineerable nucleolus in living cells by adapting a plasmid system that was previously shown to yield mature ribosomal subunits^39,43^. These rDNA plasmids contain unique sequences inserted into 18S and 28S, enabling visualization of plasmid rRNA using FISH (18S* and 28S*)^39,43^ (**Figure 3A and** **Supplementary** Figure 8A). Upon transient transfection of the rDNA plasmid, we observed formation of a *de novo* synthetic nucleolus with a characteristic FC and ring-like DFC embedded within a GC phase that recapitulates endogenous nucleolar morphology (**Figure 3B-C and** **Supplementary** Figure 10B). These synthetic nucleoli can be clearly distinguished from native nucleoli using FISH for both the unique plasmid rRNA sequences (18S*/28S*) and unique endogenous rRNA sequences - the Δ1,2,3 deleted from 5’ ETS of the plasmid (**Figure 3A-D and** **Supplementary** Figure 9A). Interestingly, these synthetic nucleoli sometimes fuse with endogenous nucleoli, generating “hybrid” nucleoli containing both endogenous and plasmid-derived rRNA (**Figure 3B-D**). We observe that 18S* plasmid rRNA localizes in discrete territories within hybrid nucleoli, suggesting that the SSU rRNA only transiently passes through the GC after processing (**Supplementary** Figure 9B), while 28S* plasmid rRNA localizes throughout the entire GC, consistent with our observations that LSU processing takes much longer in the nucleolus (specifically in the GC) (**Figure 1H**).

**Figure 3:**
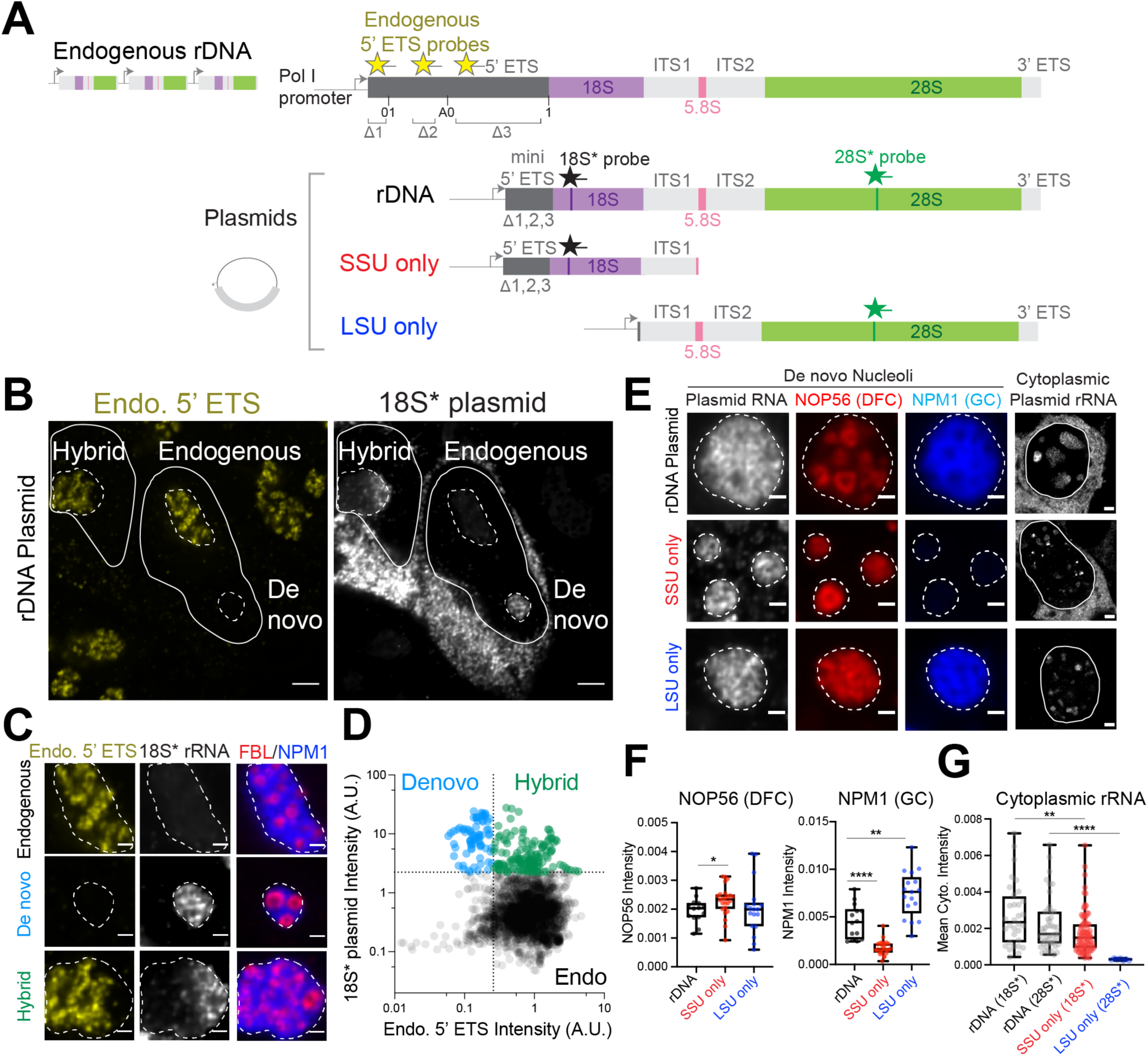
Engineered synthetic nucleoli in cells recapitulate normal multiphase architecture with the expression of the LSU precursors required for GC recruitment. **(A)** Schematics of endogenous rDNA and plasmids expressing synthetic rDNA sequences. Synthetic rDNA plasmids contain three segments (Δ1, Δ2, and Δ3) deleted from the 5’ ETS region of endogenous rDNA, enabling detection of endogenous rRNA (Endo.) 5’ ETS probes. The synthetic rDNA sequences have unique insertions in 18S and 28S that can be probed with 18S* and 28S* FISH probes. SSU only and LSU only plasmids with truncated rDNA sequences are shown. Refer to Supplementary Figure 8A-B for plasmid details. **(B-D)** RNA FISH detection of endogenous pre-rRNA by endo. 5’ ETS probes (yellow) or plasmid-expressed pre-rRNA by 18S* probe (white) in HEK293T cells after transient transfection in B. All dashed lines demarcate individual nucleoli and solid lines demarcate individual nuclei. Based on the nucleolar mean intensity of 18S* probe and endo. 5’ ETS probe, different classes of nucleoli are defined, including “De novo” (from rDNA plasmids only), “Endogenous” (from endogenous rDNA repeats), and “Hybrid” (when rDNA plasmids fuse with endogenous nucleoli), quantified in D. Example nucleoli from each class are shown in C with DFC (FBL IF) and GC (NPM1 IF). See Supplementary Figure 9A for antisense FISH probe control and Supplementary Figure 10B for de novo nucleoli visualized with all three phases labeled. **(E)** Left, de novo nucleoli formed from rDNA, SSU only, and LSU only plasmids. Plasmid RNA visualized with RNA FISH (28S* for rDNA and LSU only, 18S* for SSU only) in nucleoli labeled by DFC (NOP56-mCherry) and GC (mTagBFP2-NPM1). Right, cytoplasmic plasmid rRNA signals for each condition are shown on the side. **(F)** Quantification of mean nucleolar NOP56 (DFC) intensity and NPM1 (GC) intensity from images in E. *P value = 0.0248, **P value = 0.0012, **** P value <0.0001 (two-tailed Mann Whitney test); n = 24 (WT), 14 (SSU only), 17 (LSU only) nucleoli. **(G)** The mean cytoplasmic RNA FISH intensity for rDNA, SSU only, and LSU only plasmids quantified from images in E. ** p value = 0.0016; **** p value < 0.0001 (two-tailed Mann Whitney test). n = 32 (WT; 18S* rRNA), 47 (WT; 28S* rRNA), 90 (SSU only), 7 (LSU only) cells. Scale bars in B and E (right; whole cells) are 3 μm while the ones in C and E (left; nucleoli) are 1 μm. Box and Whisker Plots: median plotted, boxes span 25th to 75th percentiles, whiskers span min-max values.

### LSU precursors are necessary for building the GC phase

Given the spatiotemporal differences in SSU and LSU processing events reflected in these synthetic nucleoli and 5eU-seq (**Figure 1H**), we next utilized this engineerable system to ask whether SSU or LSU precursors might independently play a role in shaping distinct nucleolar phases. To do this, we generated truncated plasmids expressing only the SSU or LSU pre-rRNA sequences (**Figure 3A and** **Supplementary** Figure 8B). Remarkably, plasmids expressing only SSU precursors form ring-like DFC structures like those in native nucleoli or formed from full rDNA plasmids, but in this case there is no surrounding GC (**Figure 3E-F**). Indeed, SSU only plasmids form synthetic structures that recruit FC/DFC components (e.g., RPA194, NOP56) (**Figure 3E-F**; **Figure 4F**), but GC components are absent (e.g., NPM1, SURF6, RRP1) (**Figure 3E-F and** **Supplementary** Figure 9C), consistent with an earlier finding using a partial 18S rRNA sequence^32^. Conversely, plasmids encoding only a large subunit precursor (5.8S and 28S) give rise to a synthetic structure containing both DFC and GC components (**Figure 3E-F**), suggesting that LSU rRNA is necessary to build the GC phase. We note that 18S* rRNA expressed from the SSU only plasmid is exported successfully into the cytoplasm, further consistent with the GC being dispensable for SSU processing (**Figure 3E,G and Supplementary Note 3**). Conversely, signal for mature 28S* rRNA from the LSU only plasmid is not observed in the cytoplasm, suggesting that LSU assembly is compromised in the absence of upstream SSU pre-rRNA elements (**Figure 3E,G**). Taken together, these findings demonstrate that SSU rRNA precursors do not build the GC phase since it does not appear to be required for SSU maturation, while LSU rRNA precursors, which undergo cleavage steps in the GC, actively build the GC phase.

**Figure 4:**
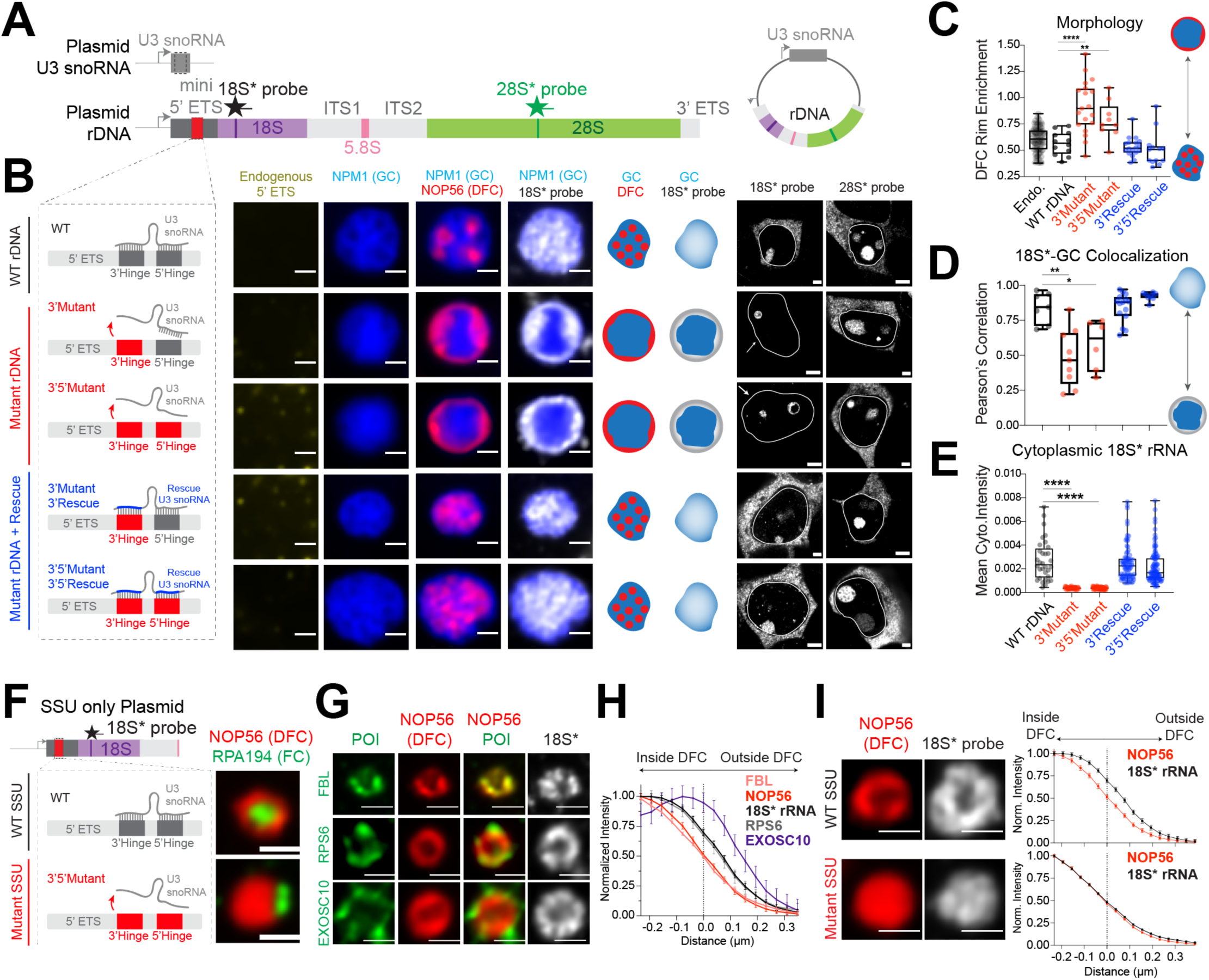
SSU processing drives the layering of the multiphase nucleolus and the outflux of SSU precursors from the nucleolus. **(A)** Schematics of plasmids expressing both U3 snoRNA and synthetic rDNA with various mutations at U3 binding sites (red). Refer to Supplementary Figure 8C-G for details. **(B)** The nucleolar morphology of de novo nucleoli as shown by the lack of endogenous 5’ ETS signal (yellow) labeled by GC (mTagBFP2-NPM1; blue) and DFC (NOP56-mCherry; red) from plasmids with normal U3 hybridization sequence, 3’ hinge mutations, 3’,5’ hinge mutations, or mutant U3 snoRNAs that rescue the hybridization. Schematics illustrate nucleolar morphology and 18S* (plasmid) rRNA localization. Cytoplasmic signals for plasmid-expressed 18S and 28S rRNAs for each condition are shown along the side. Solid lines demarcate individual nuclei. See Supplementary Figure 10B-D for all morphologies reproduced without overexpression (using IF) and visualization of all 3 nucleolar phases. **(C)** Quantification of DFC enrichment on nucleolar rim (rim enrichment score) for all plasmids in B. **** p-value < 0.0001; ** p-value = 0.0024 (two-tailed Mann-Whitney test); n = 95, 12, 19, 9, 18, 13 nucleoli **(D)** Pearson’s correlation of nucleolar 18S* signal with the GC phase (NPM1) of the nucleolus, quantified from all plasmids in B. * p-value = 0.0221; ** p-value = 0.0012 (two-tailed Mann-Whitney test); n = 7, 9, 6, 15, 8 nucleoli. **(E)** Quantification of cytoplasmic 18S* signal from B. **** p-value < 0.0001 (two-tailed Mann-Whitney test). n = 32, 38, 58, 79, 102 cells. **(F)** Schematics of wild type (WT) or mutant SSU only plasmids, refer to Supplementary Figure 8B for details, and example images of nucleoli formed by these two plasmids, labeled with DFC (NOP56-mCherry) and FC (RPA194 IF). Quantification of FC rim enrichment is in Supplementary Figure 11A. **(G)** Visualization of protein of interest (POI, shown in green), in WT SSU only nucleoli demarcated by NOP56-mcherry (red), including processing factors FBL (IF), EXOSC10 (IF), and ribosomal protein RPS6-Halotag. Plasmid-expressed 18S* RNA is shown in white. **(H)** Quantification of the radial distribution of 18S* RNA and factors in g around the DFC boundary (distance=0 is defined at 50% of maximal NOP56 signal). n = 74 (Nop56, 18S), 87 (FBL), 21 (RPS6), 12 (EXOSC10). **(I)** Examples of DFC (NOP56-mCherry) and 18S* in WT vs mutant SSU only nucleoli with their radial distribution quantified. All scale bars = 1 μm except for the right part of B (= 3 μm); n = 74 (WT SSU), 164 (Mutant SSU) nucleoli. Box and Whisker Plots: Median plotted, Boxes span 25th to 75th percentiles, Whiskers span min-max values. All error bars are s.e.m.

### Pre-rRNA processing mediated by U3 and 5’ ETS base-pairing dictates the layering of the multiphase nucleolus

Having shown that plasmid-expressed pre-rRNAs efficiently recruit assembly factors generating each nucleolar subphase (**Figure 3B-C and** **Supplementary** Figure 10B), we can further utilize this engineered system to dissect how rRNA processing shapes nucleolar morphology. Specifically, since knocking down U3 snoRNP components (i.e. U3 snoRNA or fibrillarin) results in inverted nucleoli (**Figure 2E**), we predict that designing sequence-specific changes in the plasmid rDNA to block U3-mediated processing should result in a similar morphological inversion. Briefly, we designed new plasmids containing U3 snoRNA in addition to the rDNA locus (**Figure 4A**). Using the structure of the human SSU processome as a template^39^, we mutated two sites (5’ and 3’ hinges) within the 5’ ETS rRNA spacer that directly base pair with U3 snoRNA (**Figure 4A-B and** **Supplementary** Figure 8C-G)^44–46^. We observe that the mutated rDNA plasmids form synthetic nucleoli with an inversion phenotype similar to the one observed upon U3 snoRNA knockdown (**Figure 2E**), with both FCs and DFCs localizing on the periphery of the synthetic nucleoli (**Figure 4B-C and** **Supplementary** Figure 10B-D).

Inspired by prior work in yeast^44^, we then tested if pre-rRNA processing could be restored by re-establishing base-pairing between U3 and 5’ ETS. We produced U3 variants with complementary sequences to the mutated 5’ and 3’ hinges (**Figure 4B and** **Supplementary** Figure 8C-G). Strikingly, introduction of the compensatory mutation within U3 largely restored normal nucleolar morphology (**Figure 4B-C**). This change in nucleolar structure is also correlated with restored cytoplasmic export of 18S rRNA, indicating processing is largely rescued. Specifically, mutant plasmids did not produce cytoplasmic 18S rRNA, whereas both wild-type and compensatory U3 snoRNA mutations did (**Figure 4B,E**). As a control, plasmid-expressed 28S is successfully exported to the cytoplasm in all conditions (**Figure 4B and** **Supplementary** Figure 10E).

Because the order of all three phases was inverted upon loss of U3 snoRNA function (FC>DFC>GC becomes GC>DFC>FC, see **Figure 2E**), we wanted to understand the contribution of each subphase to this global reorganization. We mutated the 5’ ETS of the minimal SSU only plasmid, which generates synthetic nucleoli lacking a GC (**Figure 3E, 4F**). While the wild-type SSU plasmid gave rise to the classical arrangement of a ring-like DFC forming around the FC (with no GC), a mutant SSU plasmid instead resulted in the DFC positioned adjacent to the FC (**Figure 4F and** **Supplementary** Figure 11A). This is similar to the peripheral localization of FCs in synthetic nucleoli produced from engineered rDNA constructs deficient for U3 binding (**Supplementary** Figure 10B-C). Taken together, these results demonstrate that base-pairing between U3 snoRNA and the 5’ ETS, a prerequisite to SSU processing, is required for the ordering and shape of the nucleolar phases.

### SSU rRNA processing is required for rRNA outflux from the DFC

Given that processing of pre-rRNA is tightly coupled with its movement through the nucleolus (**Figure 2**), we used the synthetic nucleoli to study the outflux of U3-processing deficient 18S pre-rRNA from the DFC. Unlike wild type synthetic nucleoli, where 18S rRNA precursors are localized throughout the nucleolus, U3-binding mutant 18S precursors remain localized in the DFC, and do not move into the GC (**Figure 4B,D; see schematics of RNA distribution in gray**). Compensatory U3 snoRNA mutations restore the ability of 18S precursors to be released from the DFC into the GC (**Figure 4B,D**). Thus, U3-mediated processing appears to be required for 18S pre-rRNA outflux from the DFC.

We next utilized the SSU only plasmid system (**Figure 4F**), which provides an unprecedented opportunity to study the outflux of SSU precursors without the complicating presence of the GC. Specifically, we visualized the spatial localization of 18S rRNA precursors and multiple factors demarcating various stages of SSU processing. We observed that 18S pre-rRNA and SSU processing factors are radially distributed at different distances around the DFC, with early processing factors located closest within the DFC, 18S pre-rRNA and ribosomal proteins within and beyond the DFC, and late factors at the DFC periphery (**Figure 4G-H)**. For example, early assembly factors, NOP56 and fibrillarin, components of the U3 snoRNP, are radially positioned closest to the DFC center. 18S pre-rRNA and RPS6 signal extends beyond the DFC, as RPS6 remains part of pre-40S particles after U3 snoRNP removal. Further out, EXOSC10 is localized around the periphery of the DFC, consistent with its binding upon 5’ ETS processing^39^. Altogether, our results demonstrate that components involved in distinct SSU processing steps are spatially organized within and beyond the DFC, enabling visualization of SSU outflux. These data are reminiscent of several recent proposals that there are dynamic subphases associated with the DFC in which distinct stages of processing are organized^29,47^.

This localization of 18S precursors and late SSU processing factors at the periphery of the DFC suggests that processing enables the release of RNA from the DFC phase. To test this, we utilized mutant SSU only plasmids and found that 18S precursors are no longer localized on the periphery of the DFC (**Figure 4I**). Indeed, we observe elevated intensities of 18S pre-rRNA and their early processing binding partners, ESF1, NAT10, NOP56, and fibrillarin, in SSU only mutant nucleoli (**Supplementary** Figure 11C-D), suggesting that 18S pre-rRNAs under impaired U3 processing are “trapped” with early processing factors in the DFC phase. Consistent with this accumulation of early SSU processing intermediates, the DFC grows in size in mutant SSU nucleoli (**Supplementary** Figure 11E). This lack of outflux of mutant 18S rRNA is also correlated with loss of cytoplasmic export of 18S rRNA (**Supplementary** Figure 11B). Altogether, in both the mutant SSU only and mutant rDNA nucleoli, 18S rRNA precursors appear trapped within the DFC phase and do not flux outward into the GC or nucleoplasm (**Figure 4B,D,I**).

These results suggest that the DFC phase, composed of multivalent interactions between early processing factors and pre-rRNA, can serve as a “checkpoint” to prevent the outflux of early processing intermediates. The positioning of the RNA exosome (here EXOSC10) in the periphery of the DFC, which marks the degradation of 5’ ETS and release of DFC assembly factors^39^, further underscores that processing and removal of dozens of assembly factors from SSU precursors could have an immediate effect on their ability to transition into the next compartment, namely the GC.

### An RNA-dependent multiphase model of nucleolar architecture

Our findings above indicate that irreversible steps, such as RNA cleavage and degradation events occurring during pre-ribosomal RNA processing, directly affect the morphology of the liquid-like layers of the nucleolus. As nucleoli have been described as multiphase condensates, the organization of their phases should be strongly impacted by the relative interfacial tensions (γ_i,j_) between each of the phases (i,j), which is a measure of the energy per unit area associated with interfaces^11,48–51^. The relative interfacial tensions are expected to depend on the flux and biochemical nature of the underlying molecular species^52^, including nascently transcribed pre-rRNA in the FC/DFC boundary, 18S rRNA precursors prior to 5’ ETS cleavage in the DFC, and 18S rRNA and 28S rRNA precursors in the GC (**Figure 5A**). Importantly, these rRNAs are not alone and form hundreds of interactions with proteins. Indeed, due to its interactions with many assembly proteins and RNAs, the SSU processome represents a high valency particle, whose maturation, including the cleavage and degradation of 5’ ETS, releases 50 assembly factors to generate the pre-40S particles^39,53–55^.

**Figure 5:**
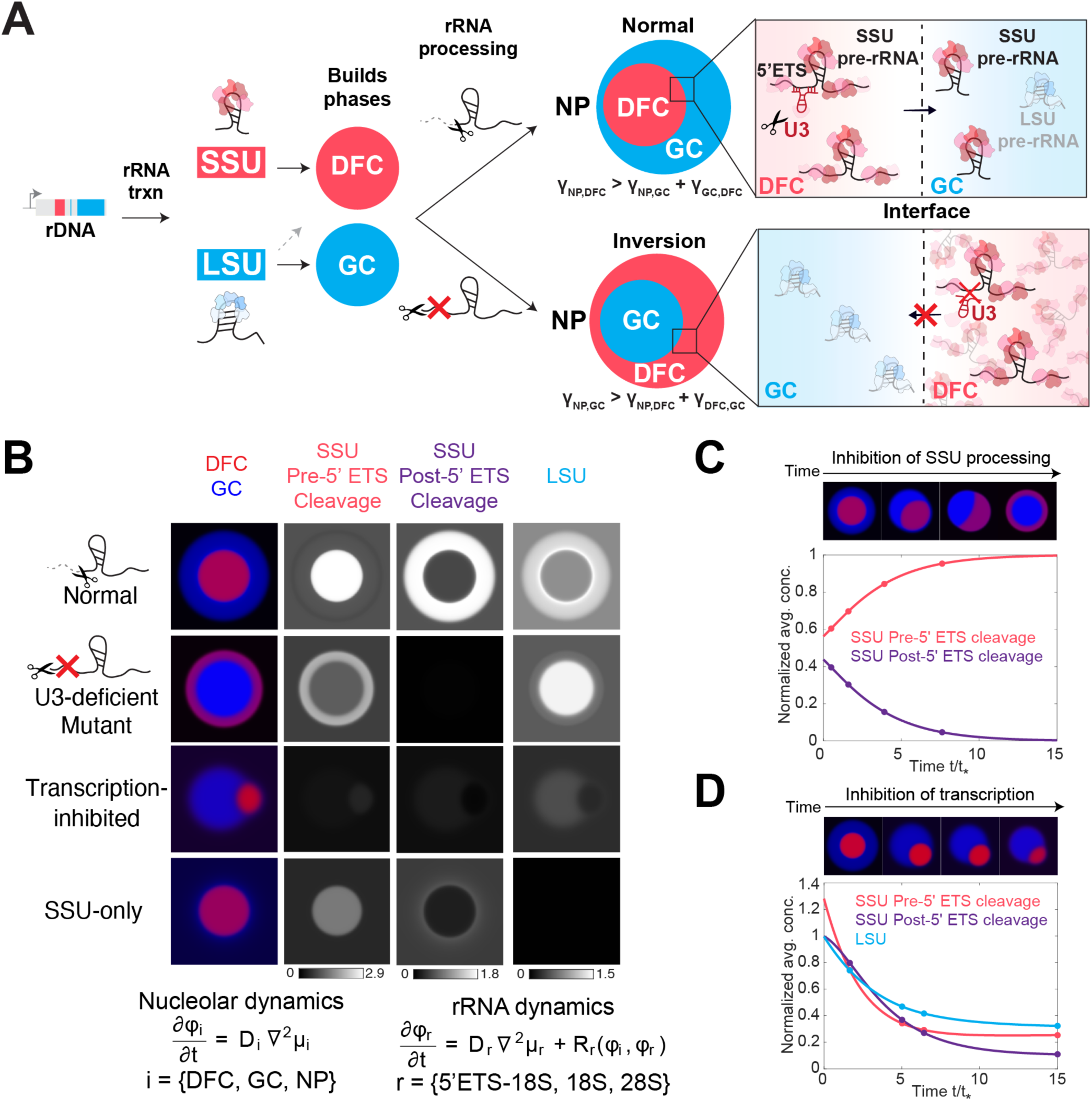
An RNA-dependent multiphase model of nucleolar architecture. **(A)** Proposed model of how rRNA transcription and rRNA processing shape the multiphase nucleolus. A 13 kb pre-rRNA is transcribed from rRNA, processed, and cleaved to assemble the SSU and LSU. Here we show that the LSU precursors are necessary for assembling the nucleolar GC phase (blue) and SSU processing drives the ordering of the DFC (red) and GC (blue) phases. Different arrangements of multiphase structures (e.g. normal or inversion) can arise from changes in interfacial tensions across multiple interfaces: Nucleoplasm (NP)-DFC (γ_NP, DFC_), NP-GC (γ_NP, GC_), GC-DFC (γ_GC, DFC_). Under normal U3-mediated cleavage of 5’ ETS from SSU pre-rRNA, the DFC localizes inside the GC. Upon impaired U3-mediated SSU processing, SSU pre-rRNAs build up in the DFC phase and are absent in the GC phase. This results in a change in the interfacial tensions and the nucleolar morphology inverts, where GC is now enveloped by the DFC. **(B)** Different nucleolar morphologies are recapitulated in a phase-field model that considers the partitioning of different rRNA precursors (e.g. SSU pre- and post-5’ ETS cleavage) into the different nucleolar phases (DFC and GC). For simplicity, the FC and DFC are modeled as one nucleolar phase. Changes in U3-mediated processing, RNA Pol I transcription, or LSU production (SSU only) alter the concentrations of rRNA precursors in each phase, resulting in different nucleolar morphologies. **(C)** Modeling of SSU processing (5’ ETS cleavage) over time whereby an accumulation of SSU precursors (pre 5’ ETS cleavage) results in inversion of the nucleolar phases. **(D)** Modeling of RNA Pol I transcriptional inhibition. Decreased concentration of all SSU and LSU rRNA precursors results in the nucleolar capping morphology.

We hypothesized that changing steady-state levels of rRNA species, namely through changes in their transcription and processing states, can impact surface tension and shape nucleolar morphology. To test this, we adapted a phase-field modeling framework well-established in material science^56^, which has also been deployed for understanding diverse condensate behaviors^3,57,58^. The system is driven to minimize thermodynamic free energy defined by the mean-field Flory-Huggins model^4,59^. Specifically, the model represents a multi-component system containing three phases, DFC, GC and nucleoplasm (for simplicity, FC and DFC were combined into a single component, namely DFC), and pre-rRNA, which is subject to production, processing, degradation, and flux consistent with the kinetic model presented earlier (**Supplementary Note 1**). A standard pairwise interaction term in the Flory-Huggins model encodes the affinity of rRNA to each nucleolar component and hence the rRNA partitioning. Specifically, given our findings that U3-processing deficient 18S precursors appear trapped within the DFC (**Figure 4B,D,I**), our model assumes that these species have high affinity for DFC, and low affinity for the GC. Conversely, because 18S pre-rRNAs with normal processing flux into GC or nucleoplasm (**Figure 4B,D,I**), we assume that 18S pre-rRNAs post 5’ ETS cleavage have low affinity for the DFC and higher affinity for the GC. In addition, we postulate that an additional three-body interaction term^60^ between rRNA and two of the nucleolar components is needed in the free energy to capture the ability of rRNA to modulate nucleolar surface tensions. Indeed, we find that this is the minimal model that can give rise to the different nucleolar morphology observed in experiments.

Under these conditions, a transition between normal nucleolar morphology and a fully inverted morphology can occur upon preventing cleavage of the 5’ ETS species, which under model parameters results in a significant increase of unprocessed species in the DFC and lack of mature 18S rRNA in pre-40S particles in the GC (**Figure 5A-B and Supplementary Note 2)**. This altered composition of the phases changes the interfacial tensions of each phase, resulting in a relocalization and final inversion of the GC and DFC phases (**Figure 5C and Supplementary Video 3**).

In addition to recapitulating the inversion morphology observed upon inhibition of SSU processing function, we tested whether our model can explain two other striking nucleolar morphologies, nucleolar “caps” classically described upon Pol I inhibition or the “SSU only phenotype” lacking a GC first reported here. Indeed, we find that reducing concentrations of all rRNA intermediates (i.e. Pol I inhibition) causes the DFCs to localize on the periphery of the GC (**Figure 5B,D and Supplementary Video 4**). Separately, the model recapitulates a strong drop in GC recruitment when the concentration of LSU intermediates are significantly reduced, much like the SSU only plasmid which lacks a GC (**Figure 5B, 3E**). While this model is a major simplification of the full complexity of pre-rRNA processing, with the details of molecular interactions implicitly encoded in the relative interaction parameters, it nonetheless clearly demonstrates a simple physical mechanism by which changes in the concentrations of rRNA processing intermediates can control nucleolar morphology through altered surface tension (**Figure 5**). Taken together, our experimental results and model demonstrate that the dynamic flux of rRNA shapes different nucleolar morphologies.

## Discussion

The nucleolus has been known for almost two centuries, but the precise origin of its multiphase architecture, and its relationship to the critical function of ribosome biogenesis, has been veiled by a lack of tools to probe this structure-function relationship. Here, we have developed and utilized a set of tools to map and dissect nucleolar organization and function, uncovering the essential components required to generate and topologically program the nucleolus. By combining spatiotemporal mapping of pre-rRNA processing, engineering synthetic nucleoli in human cells, and mathematical modeling of the determinants of multiphase nucleolar architecture, we bridge the gap between molecular scale functions of the nucleolus and its structure at the micrometer scale. Indeed, the new approaches developed here provide single-nucleotide resolution of cleavage and modification rates and allow for the parallel measurement of processing in space and time, revealing how the sequential maturation of rRNA is required for its outward flux. The ability to precisely engineer a synthetic nucleolus based on nucleotide-level changes within a recombinant rDNA locus highlights that both rRNA transcription and accurate ribosome assembly blueprint the architecture of the multiphase nucleolus.

A key finding is that although nucleolar rRNA is generated by the transcription of a single 13.3 kb pre-rRNA transcript, its cleavage, chemical modifications, and overall maturation results in SSU and LSU precursors that differentially contribute to nucleolar structure: SSU processing controls the order of the phases while the LSU is required for GC formation. Moreover, faithful pre-rRNA processing is required for the flux of pre-ribosomal particles through the successive phases, with SSU precursors being retained in the DFC if unprocessed. The nucleolar phases thus appear to represent quality control checkpoints to inhibit the release of immature precursors into the next compartment, ensuring the fidelity of ribosome assembly. Our work thus underscores the role of nucleolar multiphase organization in functional staging, where the physical separation of nucleolar phases may serve as distinct processing compartments segregating the different reactions driving the sequential maturation of SSU and LSU particles.

Altogether, we have deployed a set of cutting-edge tools to uncover how pre-rRNAs build and arrange the nucleolar phases that facilitate their processing, revealing how defects in ribosome assembly can manifest in dramatic changes to the topology of the nucleolus. Our work thus not only underscores basic principles of how RNA and its processing gives rise to the multiphase architecture of the nucleolus, but also provides a foundational toolkit for dissecting how ribosome biogenesis and nucleolar morphology are dysregulated in disease.

## Acknowledgements

We thank Y. Kang for the gifted cell lines; D. Sanders, L. Becker, A. Lin, and J. Riback for gifted plasmids; M. Guttman and P. Bhat for 5eU-seq advice; All Brangwynne lab members and H. Chang, A. Flynn, A. Herman, E. Filippova for experimental help and helpful discussions; N. Jaberi-Lashkari for manuscript feedback; E. Gatzogiannis and the Molecular Biology Confocal Imaging Facility for microscopy assistance; C. DeCoste, K. Rittenbach, G. Palmieri and the Molecular Biology Flow Cytometry Resource Facility, partially supported by the Rutgers Cancer Institute of New Jersey NCI-CCSG P30CA072720-5921, for FACS support; W. Wang and the Genomics Core Facility for genomics support.

This work was supported by the Howard Hughes Medical Institute, the Princeton Biomolecular Condensate Program, the Princeton Center for Complex Materials, a MRSEC (NSF DMR-2011750), the St. Jude Collaborative on Membraneless Organelles, and the AFOSR MURI (FA9550-20-1-0241), and the Chan Zuckerberg Initiative Exploratory Cell Network. S.A.Q. is supported by an HHMI Hanna H. Gray Fellowship. A.A.A. is supported by the Princeton University Office of Undergraduate Research, the W. Reid Pitts, Jr. Senior Thesis Fund in Molecular Biology/Biology, and the Robert W. and Eleanor A. Crecca Senior Thesis Research Fund for Molecular Biology. H.Z. is supported by the Princeton Bioengineering Institute Innovators (PBI^2^) Postdoctoral Fellowship. Q.Y. is supported by the Harold W. Dodds Fellowship from Princeton University. L.W.W. and J.F.B. are supported by the NSF GRFP Fellowship. S.K. was also supported by the National Institutes of Health (1R01GM145950 and 1R01GM143181) and the G. Harold and Leila Y. Mathers Foundation (MF-2104-01554). Research in the Lab of D.L.J.L. was supported by the Belgian Fonds de la Recherche Scientifique (F.R.S./FNRS), EOS [CD-INFLADIS], Région Wallonne (SPW EER) Win4SpinOff [RIBOGENESIS], the COST action TRANSLACORE (CA21154), the European Joint Programme on Rare Diseases (EJP-RD) RiboEurope and DBAGeneCure.

## Author contributions

S.A.Q., L.J., C.P.B., S.K. and D.L.J.L. designed the study. S.A.Q., L.J., A.A.A., L.W.W., J.F.B., A.D., E.S., L.W., and C.Z. performed experiments. S.A.Q., L.J., A.A.A., T.C., C.Z., and L.W.W. performed imaging and genomic data analysis. H.Z. and Q.Y., with advice from A.K. and C.P.B., performed simulation and kinetic modeling. S.A.Q., L.J., D.L.J.L., S.K., and C.P.B. wrote the manuscript with input from all authors. S.A.Q. and L.J. made the figures with contributions from all authors.

---

**Supplementary Figure 1:**
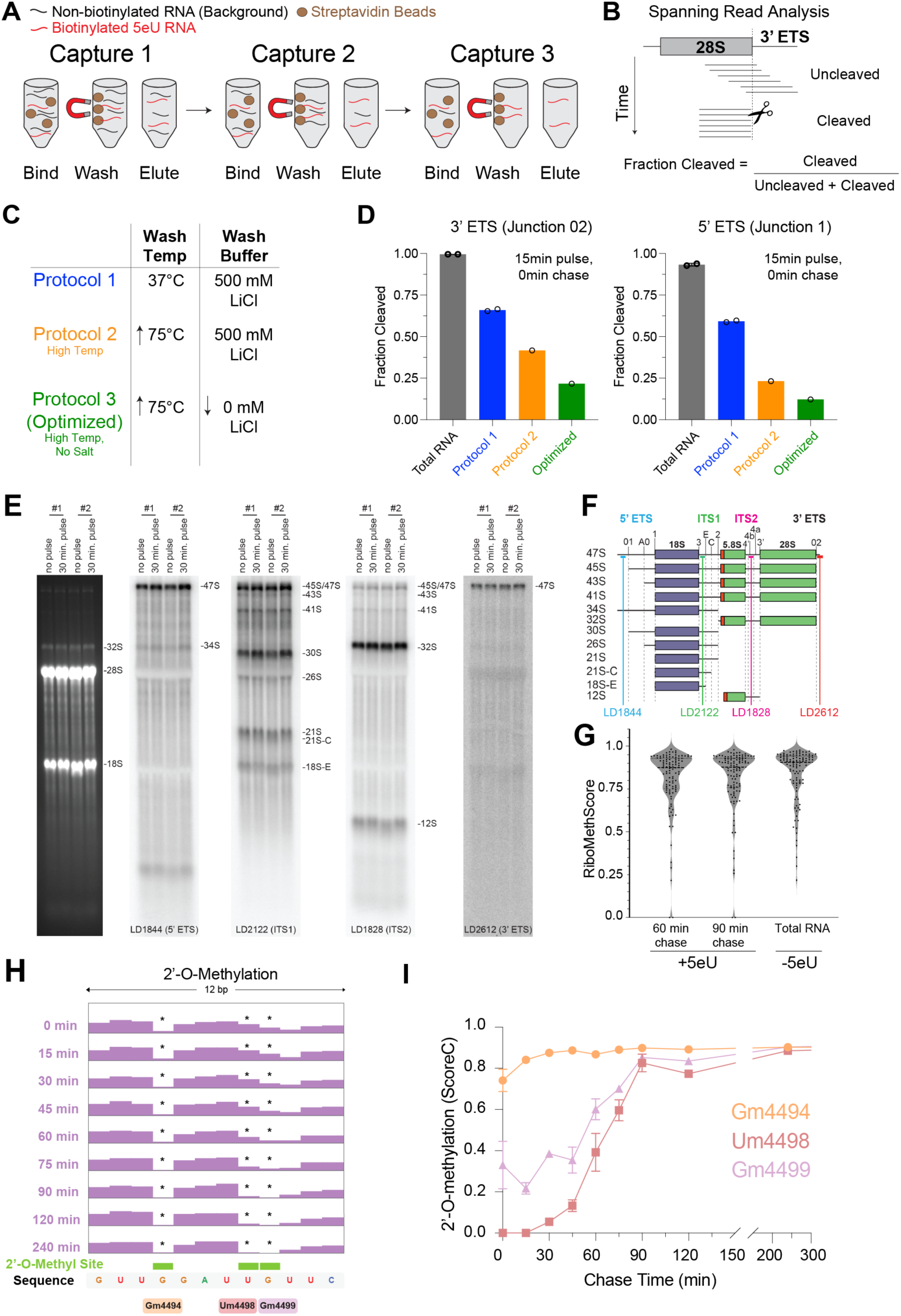
5eU-seq method description and validation. **(A)** 5eU-sequencing protocol is performed using 3 rounds of sequential captures, where biotinylated 5eU-labeled RNA is captured to magnetic streptavidin beads, washed, and eluted, as described previously^33^. **(B)** The “fraction cleaved” metric is calculated by measuring the number of “cleaved” reads, which end exactly at a cleavage junction (dashed line), divided by the total number of reads at the same position (uncleaved + cleaved). “Uncleaved” reads are reads that span a cleavage junction. **(C)** The 5eU-seq protocol^33^ (Protocol 1), which performs washes at 37° C and with 500 mM LiCl, was further optimized to reduce background from highly abundant mature rRNA, which causes the fraction cleaved metric to appear artificially high. We found that both higher temperature washes at 75° C (Protocol 2) and washing in a buffer lacking salt (0 mM LiCl) significantly reduces background (Protocol 3, Optimized). **(D)** The fraction of reads cleaved at Junctions 02 and 1. 5eU pulse-labeled material (15 min pulse, 0 min chase) from HEK293T cells was captured with streptavidin beads and washed under the Protocols 1, 2, or 3 conditions. Protocol 3 is the optimized protocol used for all datasets in this paper. Total RNA is provided as a reference for background from mature rRNA. **(E)** Northern blot analysis of 5eU pulse-labeled material (30 min) and unlabelled material as a control (No pulse). Two replicates were performed. **(F)** Northern blot probes used in E. **(G)** RiboMethScore of 2’-*O*-Methylation for 5eU 30 min pulse labeled material (+5eU, 60 or 90 min chase) and unlabelled material as a control (-5eU). **(H)** Zoom-in on 5’ end read counts of 15 min 5eU pulse-labeled material from MCF10A cells over 0-240 min chase timepoints in a region on 28S rRNA showing characteristic dips at fast (Gm4494) and slow (Um4498, Gm4499) 2’-*O*-methylation sites. **(I)** Quantification of 2’-*O*-methylation levels (ScoreC) over different chase timepoints at 28S Gm4494, Um4498, Gm4499 2’-*O*-methylation sites shown in h. Error bars are s.e.m.

**Supplementary Figure 2:**
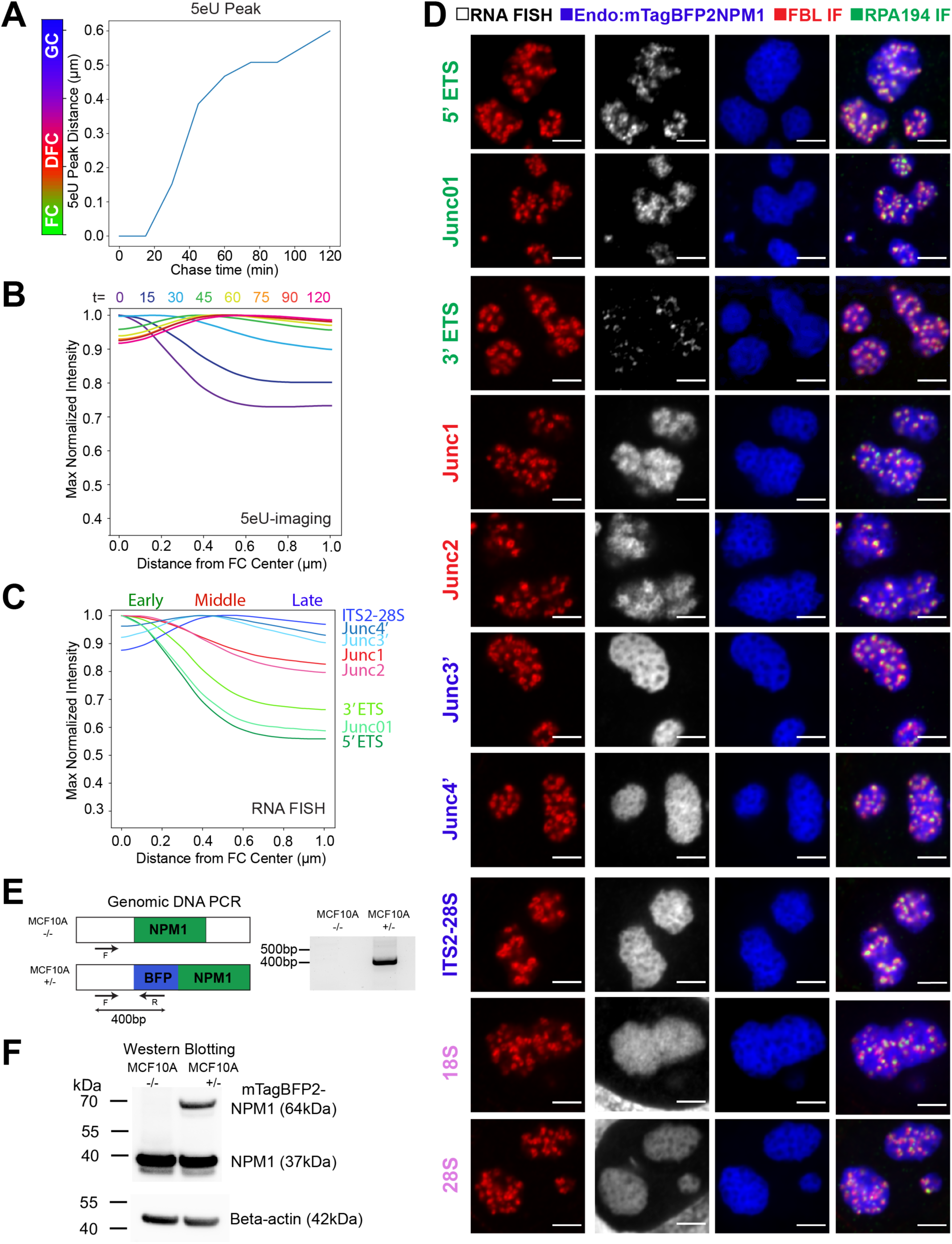
RNA FISH and 5eU-imaging of rRNA flux as well as validation of endogenously tagged mTagBFP2-NPM1 cells. **(A)** Peak of 5eU signal (distance from FC center) over chase time quantified for images in Figure 1D. **(B)** Max normalized 5eU intensity over distance from FC center over time, quantified from images in Figure 1D. **(C)** Max normalized FISH intensity over distance from FC center, quantified from images in D. **(D)** Example images of RNA FISH probes from Figure 1I-J, with FC (RPA194 IF), DFC (FBL IF), and GC (mTagBFP2-NPM1) shown. Scale bar = 3 μm. **(E)** Junction PCR of 400bp region of genomic locus spanning the inserted mTagBFP2 in MCF10A -/- (parental) and MCF10A +/-(one copy of NPM1 tagged with mTagBFP2-NPM1) cells. **(F)** Western blot for NPM1 in MCF10A -/- and MCF10A +/-cells with beta-actin as loading control.

**Supplementary Figure 3:**
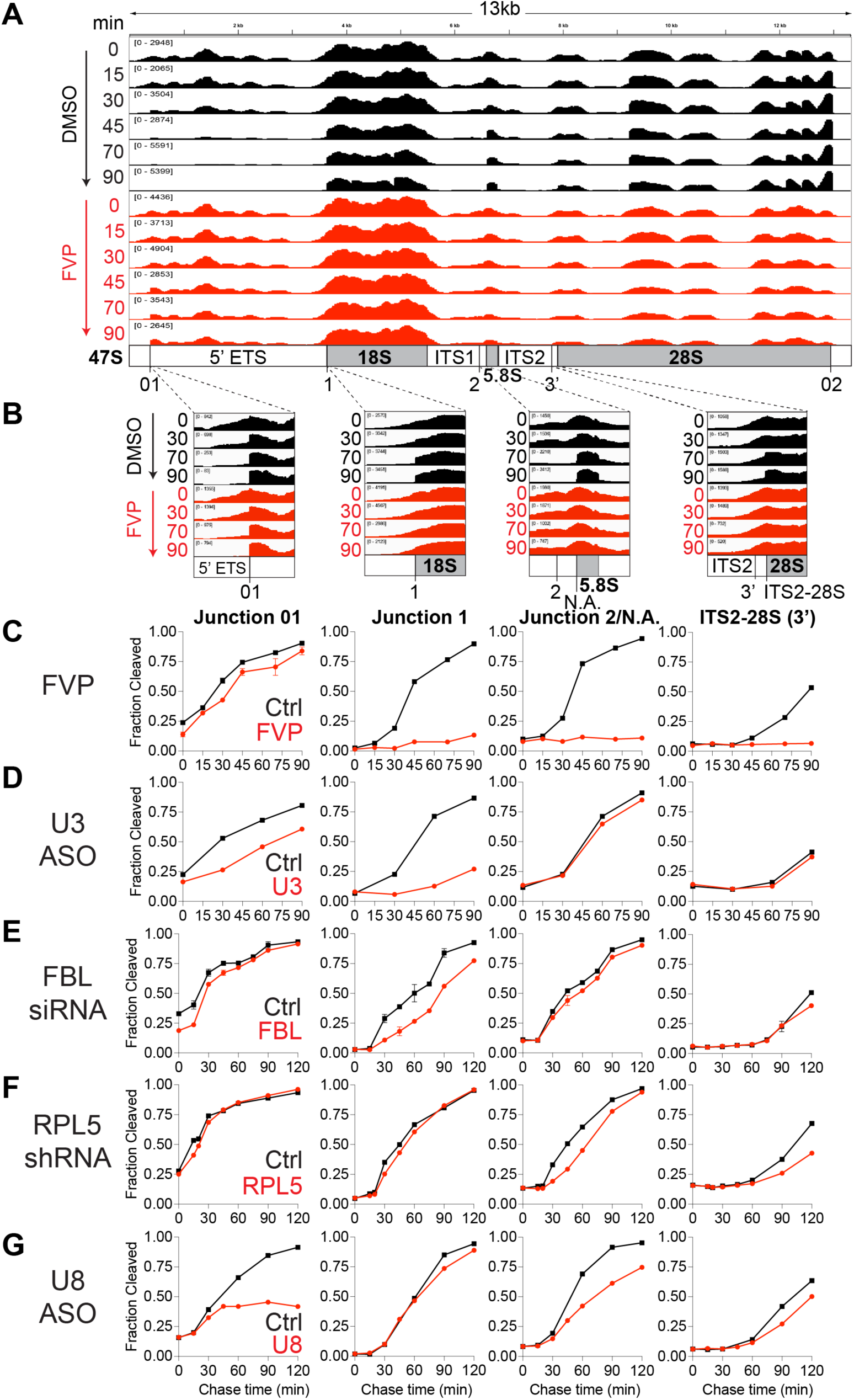
Example 5eU-seq reads and altered pre-rRNA cleavage kinetics measured by 5eU-seq upon all perturbations. **(A)** 5eU-seq reads over 47S pre-rRNA for 15 min pulse labeled material over 0-90 min chase timepoints in DMSO-treated (black) and FVP-treated (red) MCF10A cells. FVP-treated cells were pretreated with 2 µM FVP for 1 hr prior to 5eU pulse-chase and throughout the time course. **(B)** Zoom-in examples of 5eU-seq reads at 01, 1, 2/NA, and 3’/ITS2-28S regions in A. **(C-G)** Quantification of the fraction of reads cleaved at each cleavage junction displayed in B for MCF10A cells upon multiple perturbations to rRNA processing (red): FVP-treatment or knockdown of U3 snoRNA (U3 ASO), Fibrillarin (FBL siRNA), RPL5 (RPL5 shRNA), and U8 snoRNA (U8 ASO) compared to their respective controls (black; DMSO for FVP, or scramble control for ASO, siRNA and shRNA treatments). All error bars are s.e.m.

**Supplementary Figure 4:**
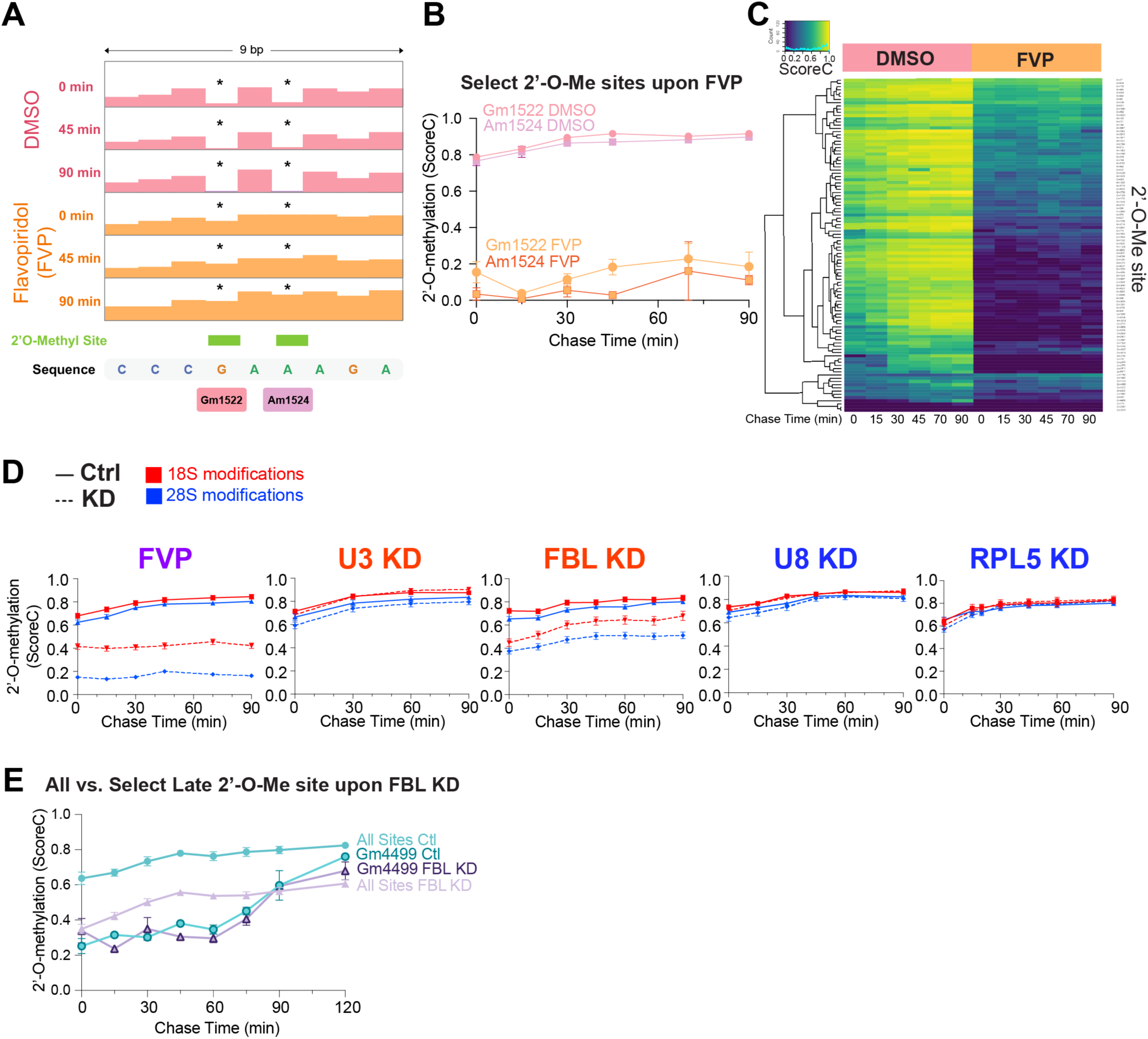
Altered pre-rRNA modification kinetics measured by 5eU-seq upon all perturbations. **(A)** Zoom-in on 5’ end read counts in control (DMSO; pink) and 2 μM FVP treatment (orange) conditions 0-90 min after transcription in 28S rRNA region at Gm1522 and Am1524 2’-*O*-methylation sites showing impairment of characteristic dips at 2’-*O*-methylation sites. **(B)** Quantification of 2’-*O*-Me levels (ScoreC) at the 28S Gm1522 and 28S Am1524 sites shown upon DMSO and FVP treatment. **(C)** Heatmap of 2’-*O*-Me levels (ScoreC) at all 18S and 28S rRNA sites in control (DMSO) and 2 μM FVP treatment 0-90 min after transcription. **(D)** Average 2’-*O*-Me levels (ScoreC) on 18S (red) and 28S (blue) rRNA in perturbations (dashed line) and control conditions (solid line) over 0 to 90 min from transcription. **(E)** 2’-*O*-Me levels (ScoreC) at 28S Gm4499 and all other 18S and 28S (average) sites in control (SCR) and FBL KD treatment conditions 0-120 min after transcription. All error bars are s.e.m.

**Supplementary Figure 5:**
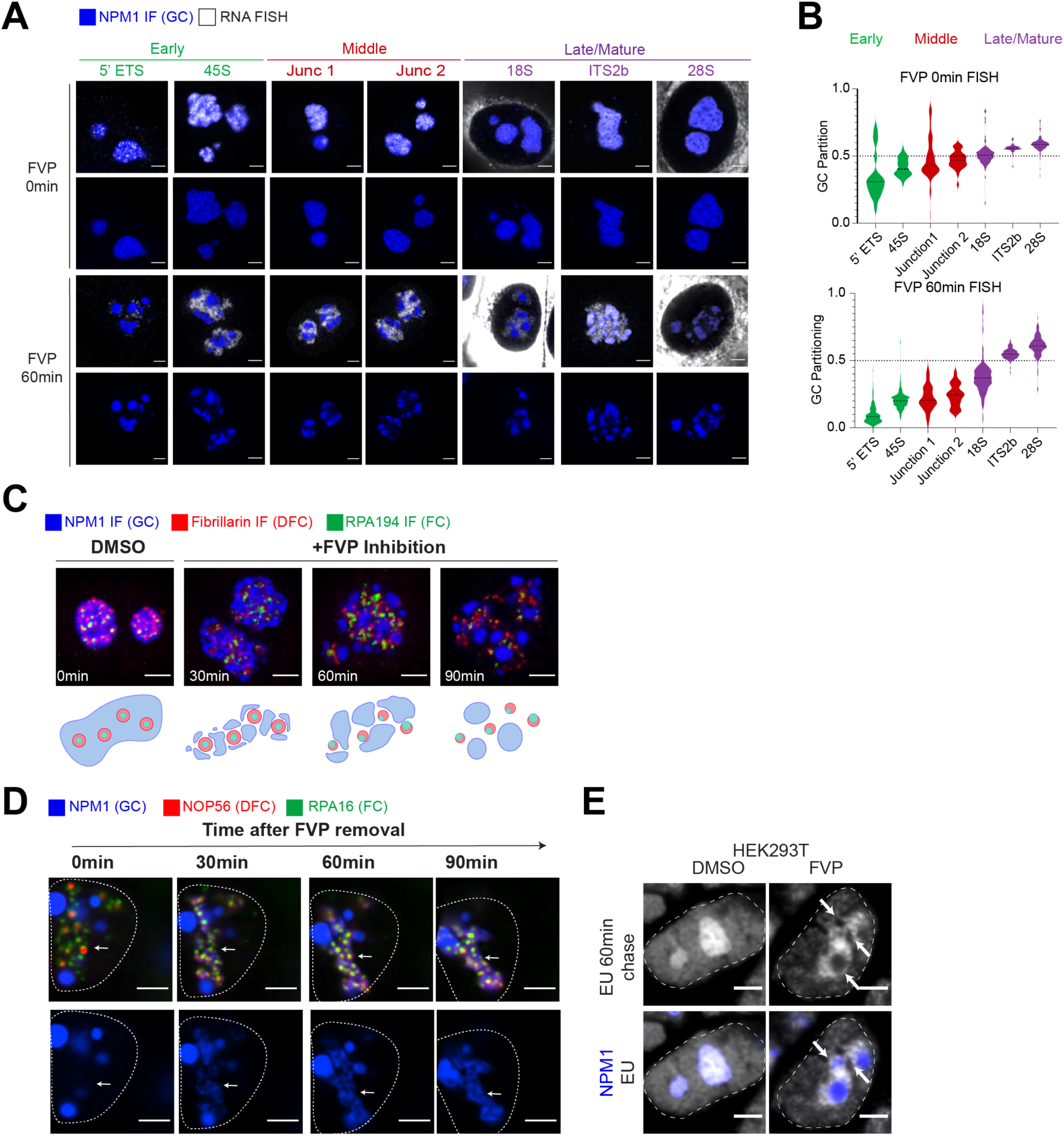
Altered localization of RNA species and nucleolar morphology upon FVP treatment. **(A)** RNA FISH for early, middle, and late rRNA cleavage sites, and mature rRNA species shown with GC (mTagBFP2-NPM1) upon 2 μM FVP treatment for 0 min and 60 min. **(B)** Quantification of GC partitioning of early, middle and late/mature RNA FISH probes upon 0 min or 60 min 2 μM FVP treatment. Violin plots are centered by median. **(C)** Nucleolar morphology showing FC (RPA194 IF), DFC (FBL IF), and GC (NPM1 IF) in DMSO and 30-90 min of 2 μM FVP treatment. Schematics show progressive GC detachment. **(D)** Nucleolar morphology showing FC (RPA16-GFP), DFC (NOP56-mCherry), and GC (mTagBFP2-NPM1) 0-90 min after wash-out of 2 μM FVP. Dashed lines demarcate nuclei. Arrows point to sites of GC reattachment to FC/DFC over time. **(E)** Example images of 5eU labeled RNA (white, 30 min pulse,60 min chase) with GC (mTagBFP2-NPM1) upon DMSO or 2 μM FVP treatment in HEK293T cells. Scale bars = 3 μm.

**Supplementary Figure 6:**
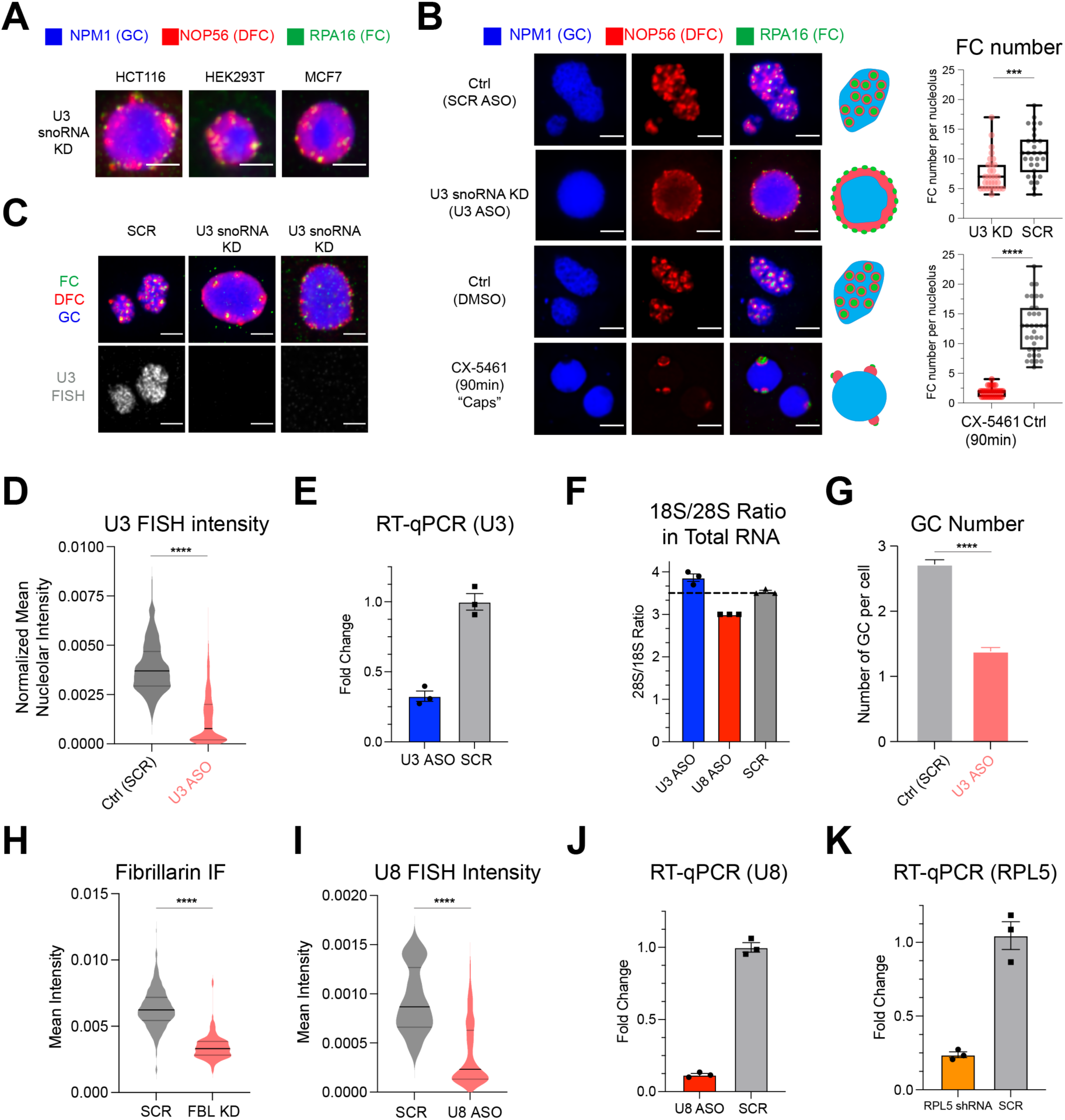
Morphology comparison between U3 snoRNA KD and CX-5461 treatment and validation of all knockdowns performed in this study. **(A)** Representative images of inverted nucleolar morphology upon U3 snoRNA KD in HCT116, HEK293T, and MCF7 cells, visualized with GC (mTagBFP2-NPM1), DFC (NOP56-mCherry) and FC (RPA16-GFP). **(B)** Left, representative images of nucleolar morphology in U3 snoRNA KD, CX-5461 treatment, and corresponding control conditions in MCF10A cells with FC, DFC, GC labeled the same as A. Right, quantification of the number of FCs per nucleolus in U3 snoRNA (n=30) and SCR (n=30) ASO (top), and CX-5461 (n=35) and control (n=31) treatment. *** p-value = 0.0002, **** p-value < 0.0001 (two-tailed Mann Whitney test). **(C)** Normal and inverted nucleolar morphology in control (SCR) and U3 snoRNA KD conditions in MCF10A cells with IF staining for NPM1 (GC), FBL (DFC) and RPA194 (FC) and RNA FISH for U3 snoRNA. **(D)** Quantification of normalized mean nucleolar intensity of U3 snoRNA FISH from C. For scramble (n=479) and U3 ASO (n=329) nucleoli. **(E)** Fold change of U3 snoRNA levels (RT-qPCR) in MCF10A cells treated with U3 snoRNA ASO and scramble (SCR) ASO for 72 hrs (n=3). **(F)** RNA electrophoresis measured 18S to 28S ratio in total RNA from MCF10A cells treated with U3 or U8 snoRNA ASO, scramble (SCR) ASO for 72 hrs (n=3 per condition). **(G)** Number of GC per cell quantified from U3 ASO (n=177) or SCR (n=384) ASO treated cells. **(H-K)** Quantification of mean nucleolar intensity of FBL (IF; n=541 SCR, 278 Fib KD) and U8 snoRNA (FISH; n=174 SCR, 40 U8 ASO), as well as fold change of U8 snoRNA (RT-qPCR) and RPL5 mRNA (RT-qPCR), respectively, under each perturbation condition (n=3 per condition). All scale bars = 3 μm. Box and Whisker Plots: median plotted, boxes span 25th to 75th percentiles, whiskers span min-max values. Violin plots are centered by median and quartiles are shown. All error bars are s.e.m. **** P-value < 0.0001 (two-tailed Mann Whitney test).

**Supplementary Figure 7:**
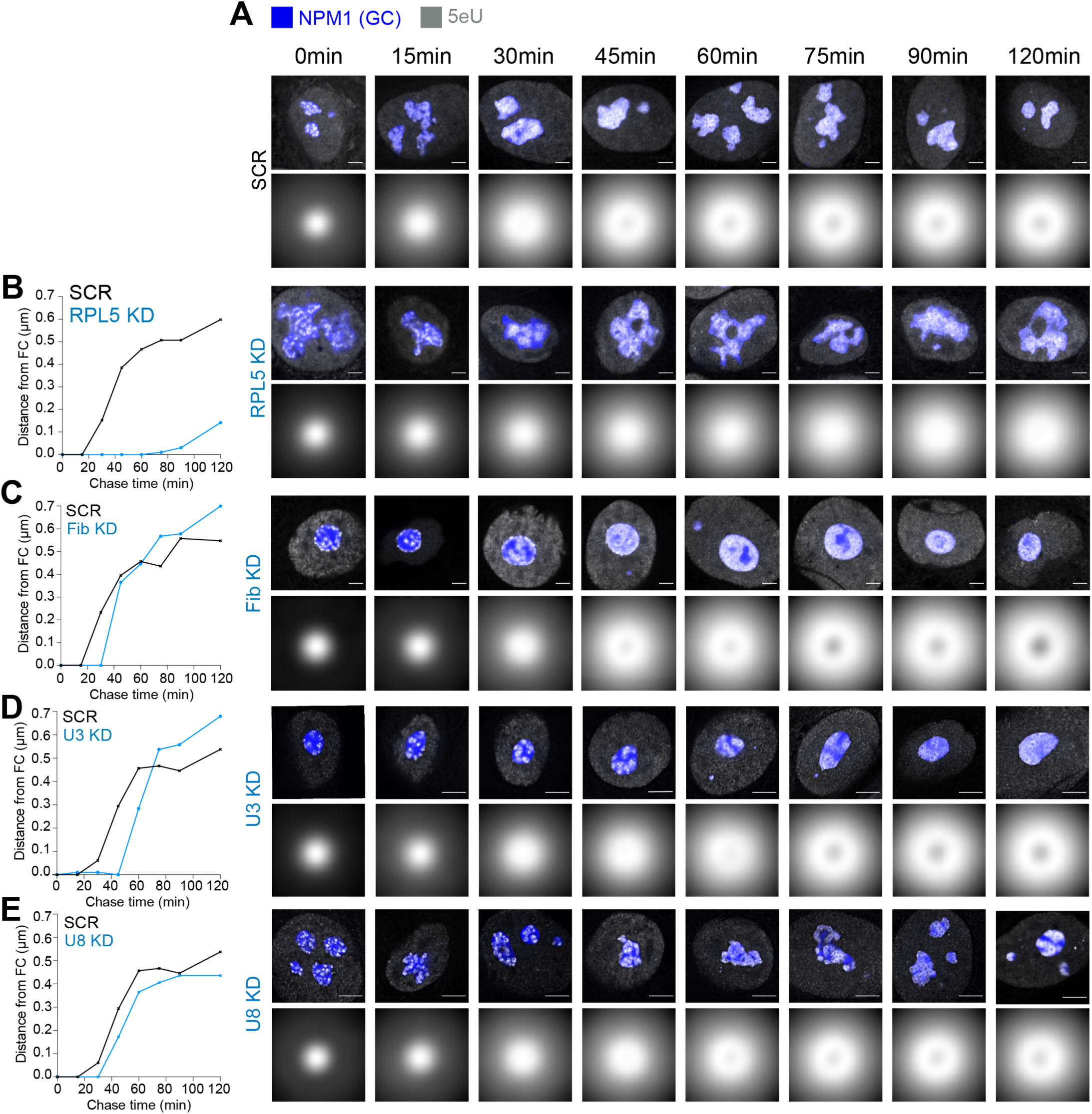
5eU imaging examples for all perturbations. **(A)** Example images of 5eU with GC (mTagBFP2-NPM1) and averaged 5eU intensity around FC in control (SCR) and all perturbation conditions over 0-120 min from transcription. Scale bars = 3 μm. **(B-E)** Quantification of 5eU peak distance from FC center over time for multiple perturbation conditions compared to corresponding controls.

**Supplementary Figure 8:**
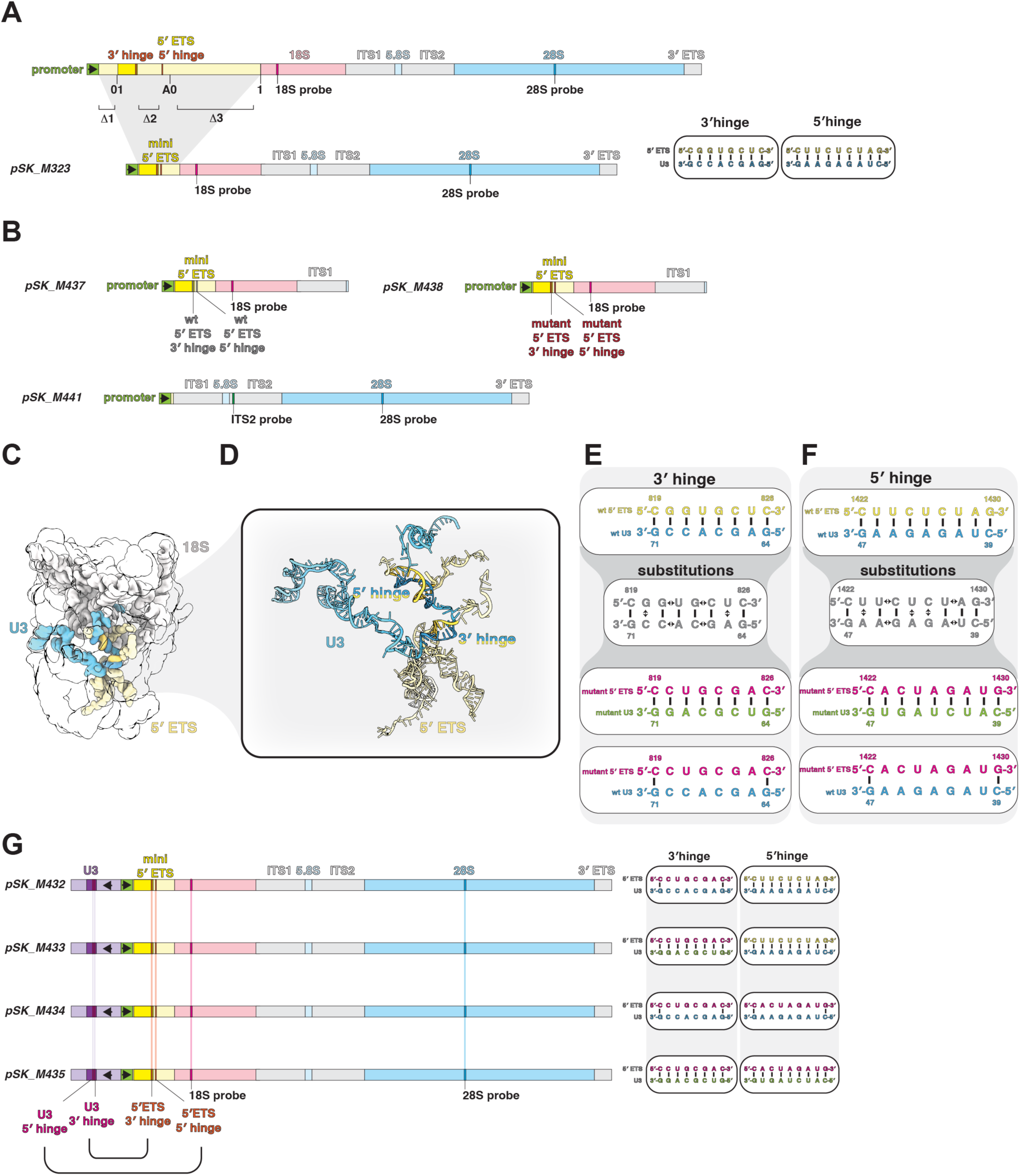
rDNA plasmid designs used in this study. **(A)** Schematics of endogenous (top) 47S rDNA and synthetic (bottom, pSK_M323) rDNA plasmid with minimized (mini) 5’ ETS. Sequences of 3’ and 5’ hinge regions of 5’ ETS-U3 snoRNA base pairing are shown. **(B)** Schematics of wildtype (WT) synthetic SSU only, 5’ ETS 3’ hinge mutant SSU only, and wildtype LSU only rDNA plasmids. **(C)** Structure of SSU processome in state pre-A1 (PDB: 7mq8). **(D)** Zoom-in on 5’ and 3’ hinge RNA duplexes between the U3 snoRNA and 5’ ETS in C. **(E-F)** Sequences of 3’ (E) and 5’ (F) hinges of 5’ ETS-U3 snoRNA base pairing in wildtype (WT), mutant 5’ ETS with mutant U3, and mutant 5’ ETS with WT U3 conditions. Sequence substitutions for mutants are marked by double-sided arrows. **(A) (G)** Schematics of complete U3 snoRNA gene combined with rDNA plasmids with 5’ ETS and U3 snoRNA 5’ hinge and 3’ hinge mutations.

**Supplementary Figure 9:**
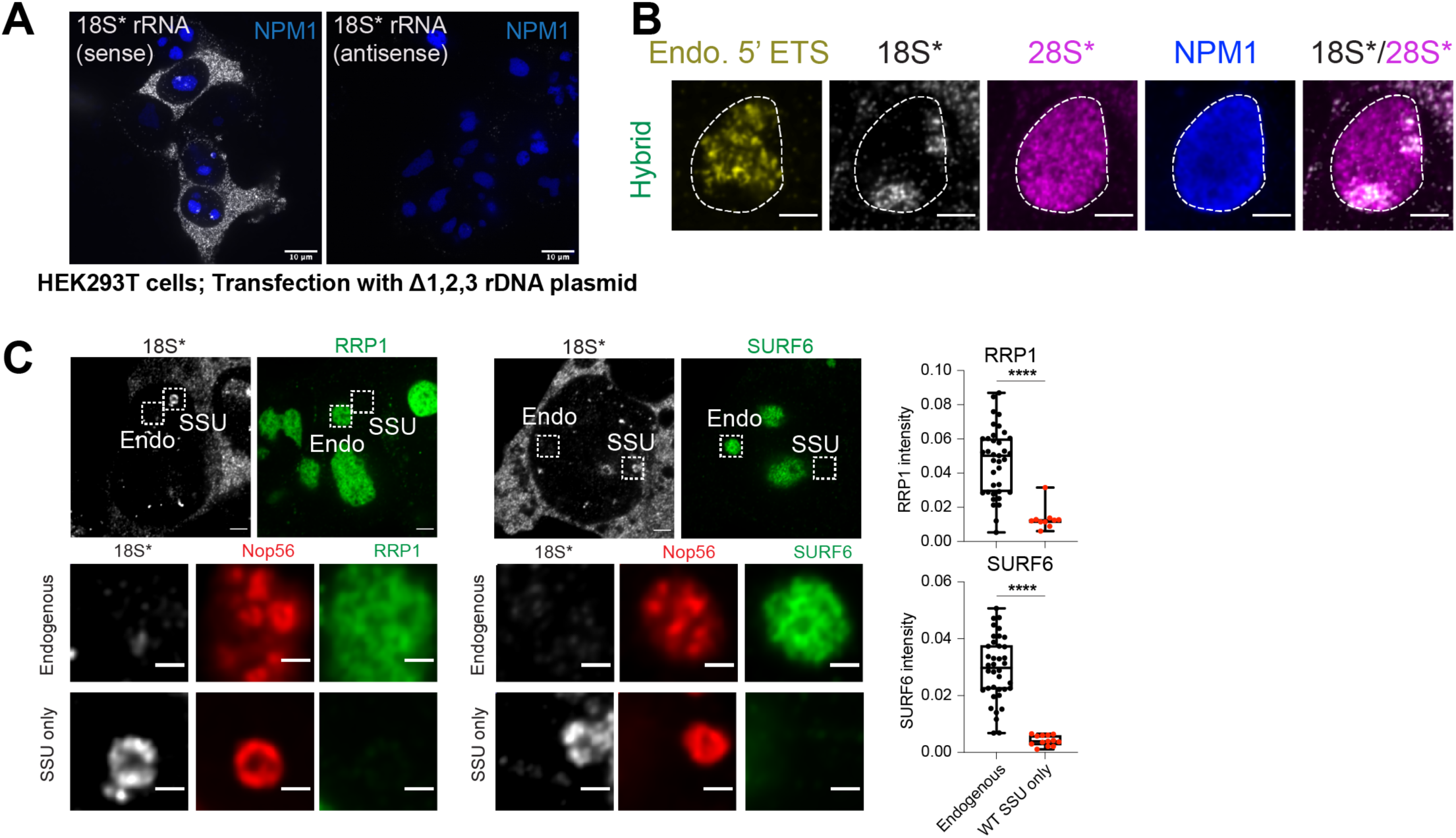
GC factors do not localize to SSU only nucleoli. **(A)** Antisense FISH probes to show RNA FISH is specific to RNA not DNA. Localization of sense and antisense 18S* RNA FISH probes with GC (mTagBFP2-NPM1) shown in HEK293T cells transfected with Δ1,2,3 rDNA plasmid. Scale bar = 10 μm. **(B)** A “Hybrid” nucleolus, as defined by signal from both endo. 5’ ETS probes and plasmid 18S*/28S* probes. 18S* labeled SSU precursors have a territory-like structure in GC (labeled with NPM1 IF) while 28S* labeled LSU precursors (transcribed from the same rDNA plasmids as 18S*) localize everywhere in GC. **(C)** Localization of RRP1 (IF) and SURF6 (IF) in endogenous and SSU only nucleoli labeled with DFC (NOP56-mCherry) and 18S* RNA FISH. Right, quantification of RRP1 (n=37, 10) and SURF6 (n=38, 13) fluorescent signal intensity in endogenous and SSU only nucleoli, respectively. Scale bars in a = 10 μm, in B and top row of C = 3 μm, in the rest of C = 1 μm. Box and Whisker Plots: median plotted, boxes span 25th to 75th percentiles, whiskers span min-max values. *** P-value < 0.0001 (two-tailed Mann-Whitney test).

**Supplementary Figure 10:**
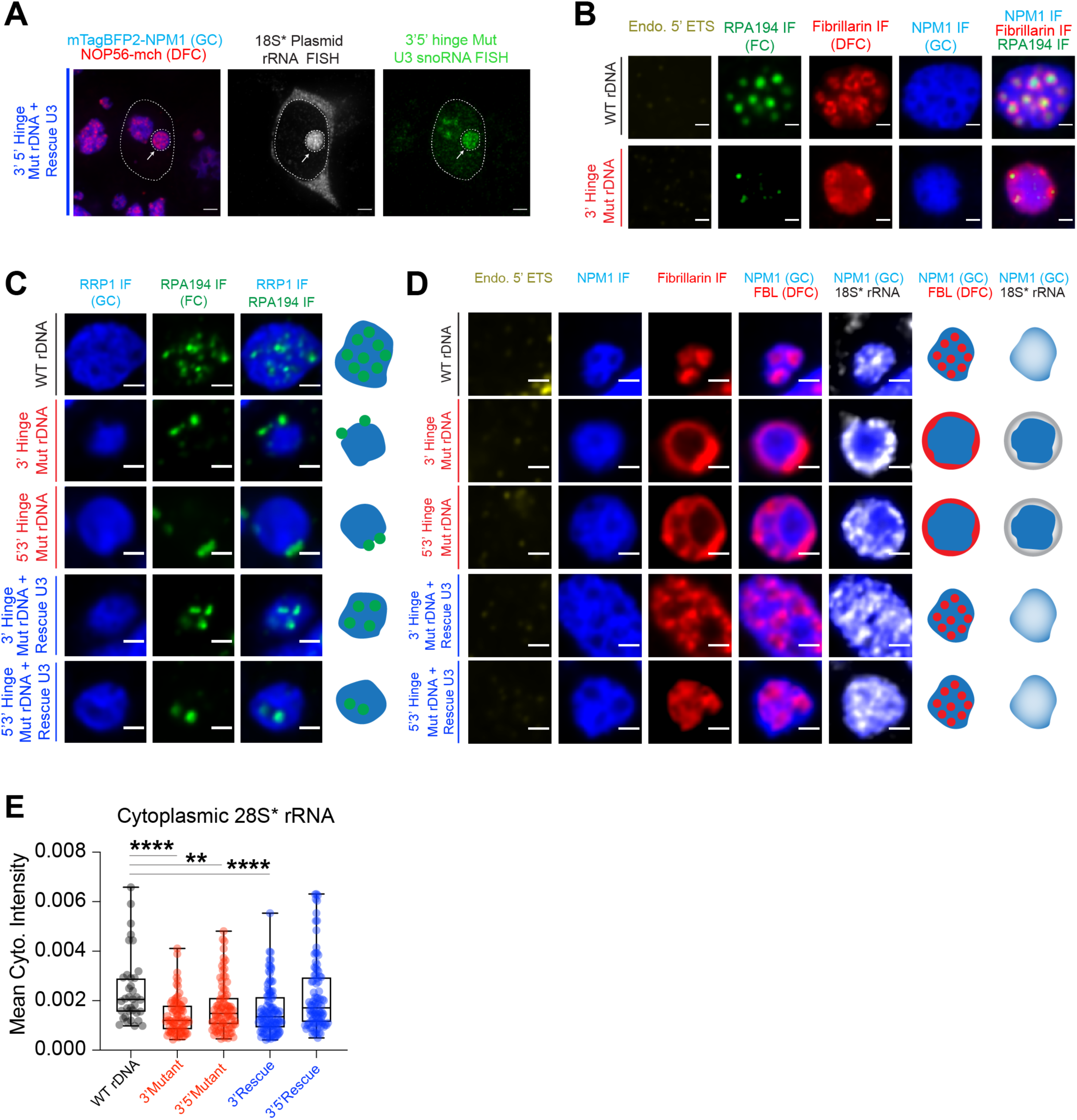
Nucleolar morphology changes upon SSU processing inhibition. **(A)** Localization of 3’ 5’ hinge mutant U3 snoRNA and 18S* plasmid rRNA by RNA FISH in cells transfected with a plasmid expressing 3’ 5’ hinge mutant rDNA and rescue U3 snoRNA. GC (mTagBFP2-NPM1) and DFC (NOP56-mCherry) are shown. Arrow points to the nucleolus with plasmid rRNA (18S*). **(B)** *De novo* nucleolus (FC labeled by RPA194 IF, DFC labeled by FBL IF, GC labeled by NPM1 IF) in cells transfected with WT or 3’ hinge mutant rDNA plasmids. Endogenous 5’ ETS is labeled by RNA FISH. **(C)** Localization of FCs (RPA194 IF) with GC (RRP1 IF) shown in cells transfected with plasmids including various mutations in U3 snoRNA binding sites. Schematics of FC and GC localization in each transfection condition. **(D)** Nucleolar morphology labeled with IF staining for DFC (FBL), GC (NPM1), and RNA FISH for 18S* rRNA and endogenous 5’ ETS in cells transfected with plasmids including various mutations in U3 snoRNA binding sites. Schematics of nucleolar morphology (DFC and GC) and 18S* rRNA localization in GC in all conditions. **(E)** Quantification of cytoplasmic plasmid 28S rRNA signal from Figure 4B. n = 36, 74, 84, 103, 92 cells. ** P-value = 0.0026; **** P-value < 0.0001 (two-tailed Mann-Whitney test). Scale bar = 3 μm for A and 1 μm for all the rest. Box and Whisker Plots: Median plotted, Boxes span 25th to 75th percentiles, Whiskers span min-max values.

**Supplementary Figure 11:**
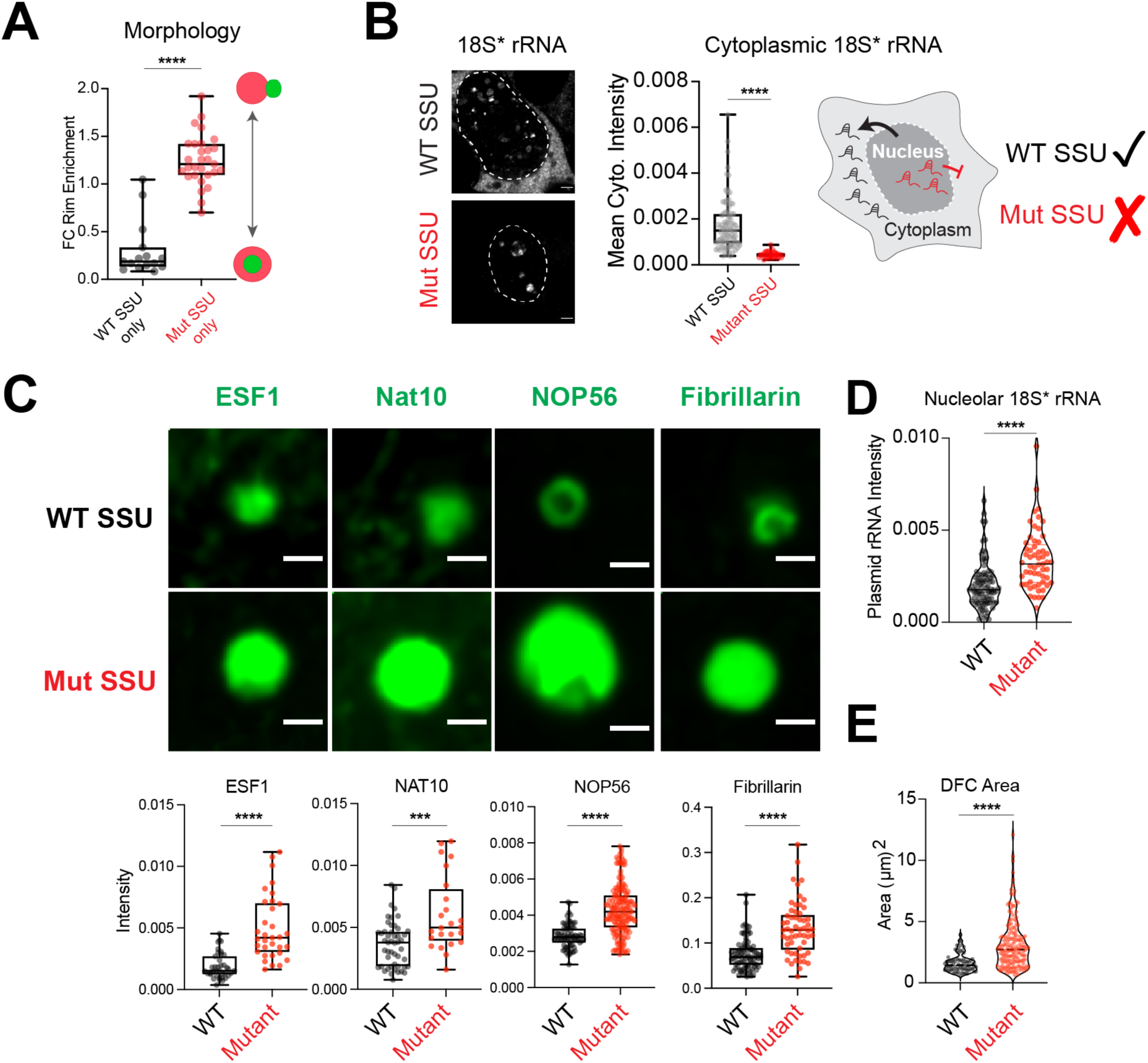
SSU only mutants have impaired cytoplasmic export and growing DFCs with increased recruitment of early SSU processing factors. **(A)** Quantification of their FC rim enrichment score from SSU WT (n=15) and SSU mutant (n=29) nucleoli shown in Figure 4F. **** p-value < 0.0001 **(B)** Localization of 18S* rRNA by RNA FISH in cells transfected with WT and mutant SSU only rDNA plasmids; 18S* rRNA mean cytoplasmic intensity is quantified in both conditions. Schematic showing normal and impaired cytoplasmic export of 18S* rRNA in WT (n=90) and mutant SSU (n=34) conditions, respectively. Scale bar = 3 μm. **** P-value < 0.0001 **(C)** Top, Localization of early SSU processing factors ESF1, NAT10, FBL (labeled by IF) and NOP56-mcherry in WT and mutant SSU conditions; bottom, mean nucleolar intensity of each is quantified in both conditions. Scale bar = 1 μm. WT SSU only n=38, 49, 74, 87; Mut SSU only n=33, 24, 164, 57 for ESF1, Nat10, Nop56, and Fib, respectively. *** P-value = 0.0005; **** P-value < 0.0001 **(D)** Quantification of mean nucleolar 18S* rRNA intensity in wildtype (WT; n=87) and mutant SSU (n=57) conditions. **** P-value < 0.0001 **(E)** DFC area (μm^2^) (labeled with NOP56-mCherry) is quantified in WT (n=74) and mutant SSU (n=164) conditions. **** P-value < 0.0001. Violin plots are centered by median. Box and Whisker Plots: Median plotted, Boxes span 25th to 75th percentiles, Whiskers span min-max values. All statistical comparisons are two-tailed Mann-Whitney tests.

## Materials & Methods

### Statistical test and sample size

All statistical tests are performed using Graphpad Prism 10 with either two-tailed Mann Whitney test or two-tailed t-tests as specified in the legends, and tests for normality were performed when appropriate. P-values and number of observations (n) are provided in the legends. For microscopy experiments, samples were prepared over multiple days (biological replicates) whenever possible and representative cells were used to create figures. For RT-qPCR measurements, technical and biological replicates were analyzed. For 5eU-seq, biological replicates were analyzed whenever possible.

### Cell culture and cell lines used in this study

All cells were cultured in a humidified chamber at 37 °C with 5% CO_2_ and 1% streptomycin and penicillin (GIBCO, 15140122). HEK293T (ATCC), HCT116 (gift from Yibin Kang) and MCF7 (gift from Yibin Kang) were cultured in DMEM (GIBCO, 11995065) supplied with 10% FBS (Atlanta Biological, S11150H). MCF10A (gift from Yibin Kang) cells are cultured in DMEM/F12 medium (Thermo Scientific, 11320082) supplied with 5% horse serum (Sigma, H1138), 20 ng/mL EGF (Novoprotein, C029), 10 ng/mL insulin (Sigma, 91077C), 1 µg/mL hydrocortisone (Sigma, H0888). For imaging, cells are treated with trypsin (Trypsin-EDTA 0.05%, Fisher Scientific 25300054) for dissociation and then seeded into 96-well glass bottom dishes (Cellvis, P96-1.5H-N) coated with bovine fibronectin (Sigma-Aldrich, F1141) diluted 1:4 in 1X DPBS (ThermoFisher, 14190144). All 5eU-seq experiments were performed on MCF10A cells.

### Recombinant human rDNA plasmids designs

A plasmid containing a previously described minimized 5′ ETS (mini 5′ ETS) that is compatible with human SSU biogenesis was used as a starting point (pSK_M323) for the design of plasmids described in this manuscript (Supplementary Figure 8A)^39^. The structure of the human SSU processome in state pre-A1 (PDB: 7mq8) was used to redesign the 3′ and 5′ hinge RNA duplexes between U3 snoRNA and the 5′ ETS (Supplementary Figure 8C-D). Nucleotide substitutions were introduced that maintain the overall nucleotide composition of the duplexes while only allowing matching variants to base pair (Supplementary Figure 8E-F). Variants of 3′ and 5′ hinges of the 5′ ETS and U3 snoRNA were combined (pSK_M432-pSK_M435) by including variants of the complete human U3 gene upstream of the RNA polymerase I promoter, resulting in a bidirectional promoter for pre-rRNA (RNA polymerase I) and U3 snoRNA (RNA polymerase II) (Supplementary Figure 8G). Plasmids coding for either wild type or mutant SSU pre-rRNAs were generated by terminating transcripts after the first 48 nucleotides of the 5.8S gene and a plasmid coding for the LSU pre-rRNAs contained the first 53 nucleotides of the mini 5′ ETS followed by ITS1, 5.8S, ITS2, 28S and the 3′ ETS (Supplementary Figure 8B). A probe for ITS2 was introduced from a previously published plasmid (pSK_M349) that was shown to give rise to mature human LSUs^43^.

### Plasmid construction

FM5-Nop56-mcherry was a kind gift from David W. Sanders. FM5-mTagBFP2-NPM1, FM5-RPA16-GFP and FM5-RPS6-Halotag lentiviral DNA plasmids were generated using the FM5 lentiviral vector (gift from David W. Sanders)^51^. A DNA fragment encoding human RPS6 was amplified from original plasmids (DNASU Plasmid Repository, HsCD00043827) by PCR with Q5® High-Fidelity 2X Master Mix (NEB) using oligonucleotides synthesized by IDT. A gene block encoding RPA16 protein was ordered from IDT. DNA fragments containing NPM1 and RPA16 were PCR amplified from gifted plasmids from Joshua A Riback. In-Fusion HD cloning kit (Takara) was used to insert the fragments into the desired linearized vector featuring a GS linker-fluorescent tag fusion. All constructs were confirmed by Sanger sequencing (GENEWIZ).

### Immunofluorescence (IF)

Cells were fixed in 96 well glass bottom plates with 4% PFA for 10 min, washed with PBS twice and then permeabilized with PBST (with 0.5% Triton X-100) for 15 min. Samples were then blocked in 2% BSA in PBS for 30 min and then incubated with primary antibodies in 2% RNAse-Free BSA (VWR, 97061-420) for 2 hrs at 37 °C (see a detailed antibody list in Supplementary Table 1). Three 1x PBS washes were conducted for 5 min each. For non-conjugated antibodies, anti-mouse or anti-rabbit secondary antibodies with the desired fluorophores are used at 1:1000 dilution for 2hrs at 37 °C. Three PBS washes were conducted with 5 min each before imaging.

| Antibody Name | Source | Identifier | Working dilution |
| --- | --- | --- | --- |
| RPA194 | Santa Cruz | sc-48385 AF488 | 1:200 |
| RPA194-488 | Santa Cruz | sc-48385 | 1:200 |
| NPM1-488 | Thermo Fisher | MA3-25200-A488 | 1:400 |
| Fibrillarin | Abcam | ab5821 | 1:300 |
| Fibrillarin-568 | Abcam | ab202540 | 1:300 |
| RRP1B | Sigma | HPA017893 | 1:100 |
| SURF6 | Thermo Fisher | PA5-54841 | 1:100 |
| Nat10 | Thermofisher | 13365-1-AP | 1:100 |
| ESF1 | proteinTech | 23496-1-AP | 1:100 |
| EXOSC10 | Sigma-Aldrich | HPA028484 | 1:100 |
| Nucleolin | ThermoFisher | 39-6400 | 1:100 |
| DDX21 | Sigma-Aldrich | HPA036593 | 1:100 |

### RNA SABER FISH

SABER FISH was performed as previously described^61,62^. Probes are designed across ribosomal RNA sequence and additional hairpin sequences are appended at the 3’ end of the probe for PER concatemerization with hairpin_28 (paired with 488 fluorescent oligos for imaging) or hairpin_25 (paired with 647 fluorescent oligos for imaging). All probe sequences are listed in Supplementary Table 1. For probing ribosomal RNAs transcribed from synthetic plasmids, 18S* and 28S* probes were designed (Supplementary Table 1) to probe the unique (∼16-20nt) insertion within the 18S and 28S sequence from plasmids^39^. Also, an antisense 18S* probe was designed to ensure the RNA FISH signal specifically comes from RNA instead of DNA (Supplementary Table 1). To specifically probe endogenous ribosomal RNA, we exploited probes hybridized to part of the 5’ ETS region that was excluded from the synthetic rDNA plasmids (Δ1,2,3)^39^. In addition, since a unique sequence is inserted within 28S (28S*), we designed a FISH probe flanking the insertion, located upstream and downstream of 28S*, for selective hybridization to endogenous 28S rRNA (endogenous 28S in Supplementary Table 1). For analyzing whether plasmid derived RNAs are exported to the cytoplasm, we used the endogenous 28S probe for segmenting the nuclei and cytoplasm (refer to “Quantitative Image Analysis” for details) IF was conducted after completing all FISH steps, starting from the blocking step as described in the IF section. Murine RNase inhibitors (NEB, M0314L) are used at 1:200 dilution in all steps of IF to preserve RNA FISH signals.

### Microscopy

A Nikon CSU-W1 SoRa spinning disc confocal equipped with the Yokogawa SoRa pixel re-assignment based super-resolution method was used for rapid super resolution imaging. The system was built around a Nikon Ti2-E fully motorized microscope and is equipped with dual Hamamatsu Fusion BT sCMOS cameras. The W1 Sora system is equipped with 405, 488, 514, 532, 561, 594, and 640 nm laser lines. For this work, a Nikon CFI Plan Apo Lambda D 60x oil (MRD71670) was used with a 2.8x SoRa magnification and 405, 488, 561, 640 nm lasers. A Mad City Labs piezo z stage was used for z-stack acquisition. All acquisitions were performed with the Fusion BT in “Ultra Quiet” readout using correlated double sampling. In some cases, Denoise.ai (Nikon software) was performed for images shown and analyzed in this study.

### Lentiviral packaging and transduction

Lentiviruses were made using HEK293T cells seeded in a 6 well plate at 70-80% confluence. The desired plasmids were transfected together with helper plasmids VSVG and PSP via lipofectamine^TM^ 3000 (Invitrogen, L3000008) following previously described protocols^51,63^. Viruses were collected 48 hours after transfection and filtered through syringe filters with 0.45 µm pore size (VWR). Lentiviral transduction was conducted in 96-well plates at 30% cell confluency for 2 days and then cells were expanded to make stable expression lines and FACS sorted for polyclonal lines with a tight window for each fluorescent protein intensity.

### Transient Transfection

rDNA plasmids (including engineered mutant plasmids) were transfected into HEK293T cells (for one 24 well, 600 ng plasmids are transfected with 150K HEK293T cells) using lipofectamine^TM^ 3000 (Invitrogen). Cells were seeded into 96 well glass bottom plate 24hrs post transfection and fixed for SABER FISH 48 hrs post transfection. For SSU only plasmids, cells were imaged 24 hrs post transfection.

### Endogenous N-terminal tagging of NPM1 with mTagBFP2 using CRISPR/Cas9

Endogenous N-terminal tagging of NPM1 in MCF10A cells was done as previously described^64,65^. An oligonucleotide pair encoding an NPM1-targeting gRNA (TGTCCATCGAATCTTCCAT) was cloned into a modified lentiCRISPRv2-puro plasmid (courtesy of Aaron Lin) via the BsmBI restriction site. MCF10A cells were transfected using the FuGENE® HD Transfection Reagent (Promega, E2311) with plasmids containing the cloned gRNA and a donor plasmid. The donor plasmid was constructed by cloning the tag with a flexible linker flanked by 300 bp homology arms complementary to the N-terminus of the NPM1 gene into the pUC19 vector (ThermoFisher, SD0061). Three days after transfection, mTagBFP2-positive cells were single-cell sorted into 96-well plates. These single cell clones were then cultured and expanded for tagging validation through western blotting, junction PCR of the specific genomic locus (Supplementary Figure 2E-F), and imaging to confirm correct subcellular localization.

### All ribosomal RNA processing perturbations used in this study

*CX-5461 Pol I Inhibition:* Cells were treated with 10 μM final CX-5461 (MedChem Express; HY-13323) for 90 min before fixation and imaging. DMSO was used for the control group.

*Flavopiridol Broad rRNA processing Inhibition:* Cells were treated with Flavopiridol (FVP) at 2 μM final concentration (MedChem Express; HY-10005). As a control, cells were treated with DMSO. For the 5eU-imaging and 5eU-sequencing, cells were pretreated for 1 hr to broadly inhibit processing prior to a 15 min pulse and subsequent 0-90 min chase. All pulse-chase media contained 2 µM FVP to ensure processing inhibition was maintained throughout the 5eU pulse-chase labeling. For imaging the reformation of the multiphase nucleolus upon FVP washout, cells were treated with 2 μM FVP for 2 hrs and then washed twice quickly with 1X DPBS before imaging in regular medium. Movies were taken at 37 °C with 5% CO2, every two minutes after FVP removal over 90 min.

*RPL5 shRNA*: For shRNA vector cloning into a lentiviral plasmid for expressing shRNAs with puromycin selection, RPL5 shRNA (GATGATAGTTCGTGTGACAAA) sequences and a negative control (GCTCTTAACTAACGTCACCTA) sequence were separately cloned into pLKO.1 TRC vector after digestion with AgeI and EcoRI. Lentiviruses were produced as described above and added to cells for 5eU-sequencing and imaging experiments with 30% confluency of MCF10A cells. After one day, virus-containing medium was removed and replaced with fresh medium including 5 µg/mL of puromycin. After selection for another 3-4 days, cells in the negative control well not treated with virus were dying, indicating selection was effective, and selection was terminated by replacing the puromycin medium with normal MCF10A medium. Cells were then split for 5eU sequencing, imaging or RNA extraction followed by RT-qPCR (Supplementary Table 1) 4-5 days post adding shRNA viruses.

*U3/U8 snoRNA ASO*: U3 and U8 snoRNA ASO treatment was performed as previously described^37^ with several adaptations. For 5eU-seq experiments, in each well of a 6 well plate, 1.5 μL of 40 μM (stock) ASO diluted in 125 μL Opti-MEM (ThermoFisher, 31985062) was combined with 7.5 μL Lipofectamine RNAiMAX (ThermoFisher, 13778075) diluted in 125 μL Opti-MEM. After a 30 min incubation at room temperature (RT), 1.75 mL of a suspension containing 250,000 cells in antibiotic-free medium (refer to ‘Cell cultures and lines used’) was added to each well. Cells were incubated for 1.5 days prior to performing 5eU-seq. For imaging experiments: In each well of a 96 well plate coated with bovine fibronectin (Sigma-Aldrich, F1141) diluted 1:4 in 1X DPBS (ThermoFisher, 14190144), 0.05 μL of 40 μM ASO diluted in 4.165 μL Opti-MEM was combined with 4.165 μL Lipofectamine RNAiMAX diluted in 0.25 μL Opti-MEM. After a 30 min incubation at RT, 100 μL of a suspension containing 8,000 cells in antibiotic-free medium was added to each well. For 5eU-imaging experiments, cells were incubated for 3 days prior to fixation. For experiments monitoring the inversion morphology, cells were treated again with ASO after 2.5 days as per the protocol above, then incubated for an additional 2 days prior to fixation. All incubations were performed at 37 °C and 5% CO_2_. ASO sequences were published in a previous paper^37^ and are listed in Supplementary Table 1.

*Fibrillarin siRNA*: Fibrillarin siRNA treatment was performed as described in the U3 and U8 snoRNA ASO treatment protocol with the following modifications. For 5eU-seq experiments and western blotting, 2 μL of 20 μM fibrillarin siRNA (Supplementary Table 1) or control siRNA (ThermoFisher, 4390843) diluted in 250 μL Opti-MEM was combined with 6 μL Lipofectamine RNAiMAX diluted in 250 μL Opti-MEM. For imaging experiments: 0.066 μL of 20 μM fibrillarin siRNA or control siRNA diluted in 8.33 μL Opti-MEM was combined with 8.33 μL Lipofectamine RNAiMAX diluted in 0.2 μL Opti-MEM. For all experiments, cells were treated again after 24 hours with fibrillarin siRNA or control siRNA. Cells were incubated for an additional 3 days prior to performing 5eU-seq, harvesting for western blotting, or fixation for 5eU-imaging.

### Total RNA isolation

For each well of a 6 well plate, total RNA was harvested in 200 μL 1X Buffer RLT (QIAGEN, 79216) and isolated using the QIAGEN RNeasy Mini Kit (74104), followed by 1 hr of DNase treatment using TURBO DNase (ThermoFisher, AM2238). DNase-digested RNA was then further purified with the Zymo RNA Clean and Concentrator-25 kit (R1017).

### RNA electrophoresis

RNA integrity post-isolation was analyzed and the ratios of 28S to 18S rRNA in U3 and U8 snoRNA ASO and SCR ASO-treated samples were assayed using the Agilent RNA High Sensitivity Assay (Agilent, 5067-5579) on the 4150 TapeStation system (Agilent Technologies, Santa Clara, CA, USA) as per the manufacturer’s instructions.

### qPCR for validation of perturbation

U3 snoRNA, U8 snoRNA, and RPL5 mRNA knockdown efficiency were assayed by real-time quantitative reverse-transcription PCR (RT-qPCR) using the Luna Universal One-Step RT-qPCR Kit (NEB, E3005) as per the manufacturer’s instructions, except that each reaction was scaled to 60 μL to allow for 4 technical replicates (12.5 μL) per sample. U6 snoRNA served as a loading control. RT-qPCR was performed on an Applied Biosystems QuantStudio 3 Real-Time PCR System instrument (A28567). Primer sequences used are listed in Supplementary Table 1. All primers were synthesized by Integrated DNA Technologies (IDT). U3, U8, and U6 snoRNA absolute amounts were determined using standard curves prepared for each primer set. U3 and U8 knockdown efficiency was assayed by comparing the absolute amounts of U3 or U8 snoRNA normalized to absolute amounts of U6 snoRNA, in U3 or U8 snoRNA ASO versus SCR ASO-treated samples.

### 5eU labeling combined with imaging (5eU-imaging)

5eU-imaging protocol was modified from previous studies^34^. Briefly, cells were seeded a day before at 40% confluency in 96 well glass bottom plates. Volumes of all reagents were kept at 100 μl for each 96 well. 5eU (Thermofisher, E10345) solution was prepared at 0.5 mM in cell culture medium and added to cells with medium removed from the well. Then cells were kept in an incubator (37 °C with 5% CO_2_) for 15 min (this is called “pulse”). Next, 5eU containing medium was removed and quickly washed twice with 1x DPBS containing 10 mM (excess) uridine (Sigma, U6381-5G) to outcompete the leftover 5eU. Then, culture medium containing 10 mM uridine was added to the well before incubating for different chase time points (0 min to 120 min) at 37 °C with 5% CO_2_. Note that all solutions mentioned above were kept at a 37 °C heat block to minimize temperature induced effects on RNA transcription and processing. For wells with different chase times on the same 96-well plate, the starting time of the 5eU pulse was staggered so that all the chase time points end at the same time. For fixation, 4% formaldehyde (PFA) diluted from 16% PFA (Fisher Scientific, PI28906) with 1x PBS was used for 15 min at RT. Fixed cells were then washed with 1x PBS twice and permeabilized with 1x PBST (1x PBS; 0.5% Triton X-100) for 15 min. For click chemistry, Click-iT™ Plus Alexa Fluor™ 647 Picolyl Azide Toolkit (Thermo Fisher, C10643) was used following manufacturer’s instructions, with the exception of using AZDye 647 Picolyl Azide (Click chemistry tools; CCT-1300-1) instead of the azide from the kit. After 30 min of applying click chemistry reaction cocktail, cells were washed once with 1x PBS before imaging. For combining IF with 5eU imaging, IF steps were performed as described above before click chemistry.

### 5eU labeling combined with sequencing (5eU-seq)

All 5eU pulse-chase labeling experiments were performed in MCF10A cells except for method validation described in Supplementary Figure 1, which was performed on HEK293T cells. Cells were seeded for 5eU pulse chase labeling such that their confluence was ∼70-80% at the time of harvesting. Cells were pulse labeled with 5eU (Jena Biosciences; CLK-N002-10) for 15 minutes at 37 °C with 5% CO2. Cells were then removed from the incubator and quickly washed twice with 1x DPBS to remove 5eU from the cells. Cells were then chased with media containing 10 mM Uridine (Sigma, U6381) to outcompete the leftover 5eU in cells over different chase time points. All solutions were kept at 37 °C using a heat block to minimize temperature-induced changes to the cells, which could impact RNA transcription and processing. After the given chase time, cells were harvested in 1x Buffer RLT (QIAGEN, 79216) and frozen at -80° C for RNA isolation. RNA isolation was performed with Qiagen RNeasy kits followed by DNAse digestion to remove genomic DNA. RNA concentrations were measured with Qubit RNA BR (Fisher Scientific, Q10211) and RNA integrity was analyzed on RNA HS Tapestation.

Total RNA (10-15 µg) isolated after 5eU-pulse chase was click-reacted with biotin picolyl azide (Click chemistry tools, 1167-25) as described^33^ with the following modifications. Capture of biotinylated RNA was performed using 20uL Dynabeads MyOne Streptavidin C1 beads (Invitrogen, 65002) after click chemistry. Captures and washes were performed as previously described with the following modification. The 3 x 5 min washes of captured material were changed to 75° C in No Salt Urea buffer (4M Urea, 10mM HEPES, pH 7.5, 10mM EDTA, 0.5% Triton X-100, 0.2% SDS, 0.1% Na-DOC). We found that 3 rounds of sequential captures (as described by Bhat et al, 2024^33^) as well an optimized protocol introduced here performing washes at high temperature and in a buffer lacking salt were all essential to reduce background of highly abundant mature rRNA (Supplementary Figure 1A-D). RNA-seq library prep was performed as previously described^66^. 5eU-seq libraries were then sequenced on a NovaSeq 6000 (Illumina) with paired-end reads (either 150x150 or 100x200).

### High performance computing

The analyses presented in this article were performed on computational resources managed and supported by Princeton Research Computing, a consortium of groups including the Princeton Institute for Computational Science and Engineering (PICSciE).

### Computational analysis of 5eU-seq cleavage and 2’-*O*-methylation (5eU-seq)

A custom snakemake pipeline was used to perform alignments, 2’-*O*-methylation analysis, and cleavage analysis: https://github.com/SoftLivingMatter/5eU-seq-pipelines. Sequencing reads were trimmed with Trimmomatic v0.39^67^ to remove adaptor sequences and bases containing low quality scores. Trimmed reads were then aligned to the rDNA genome (GenBank: U13369.1) using STAR (2.7.11a)^68^. Reads were sorted and indexed using Samtools (v1.9-4)^69^ and only uniquely mapped reads were kept for further analysis. The fraction of reads that are cleaved at a given junction was calculated as the number of non-spanning reads divided by the total number of reads at a given junction (Supplementary Figure 1A) using featureCounts v1.6.4^70^ (Subread package). rRNA cleavage junctions were annotated based on positions described in Mullineux and Lafontaine 2012.^15^ All rRNA junctions used in this study were manually inspected to define the sites of cleavage based on where the 5’ or 3’ end of reads piled up, demarcating a precise cleavage site. In cases where rRNA cleavage events are followed by gradual degradation, a broader window downstream of an annotated junction was used.

To determine 18S and 28S ribosomal RNA 2’-*O*-methylation levels over time, we applied the RiboMeth-seq computational analysis^71,72^ to 5eU-seq sequencing reads. We calculated ScoreC at known 2’-*O*-methylation sites ^41^ using a weighted average of the 5’ end read counts in a +/-2 nucleotide window, recommended by Pichot et al. 2020^73^, around each site. Nucleotides in the +/-1 and +/-2 neighboring positions were assigned weight contributions of 0.9 and 1, respectively. If a separate 2’-*O*-methylation site was found within the +/-2 window around a site, the 5’ end read counts at the former were skipped and those at the immediately preceding nucleotide were used alternatively. For the calculation of ScoreC, nucleotide positions of 2’-*O*-methylation sites on 18S and 28S rRNA were converted to those on 47S rRNA by adding +3655 to the 18S 2’-*O*-methylation positions (except Am1678, Cm1703, Um1804, to which +3657 was added) and +7903 to 28S 2’-*O*-methylation positions. For data visualization, negative ScoreC values were clipped to zero. The heatmap in Supplementary Figure 4C was generated in R using the heatmap.2 function from the gplots package.

### RiboMethSeq

The RiboMethSeq was conducted exactly as described in Marchand et al. 2016^71^ for data in Supplementary Figure 1G. *Analysis*: Adapter sequences were trimmed from raw reads using Trimmomatic v0.39 with the following parameters: LEADING:30, TRAILING:30, SLIDINGWINDOW:4:15, AVGQUAL:30, and MINLEN:17. Quality control of the raw and trimmed reads was assessed using FastQC (v0.11.9). Alignment to the reference rRNA sequence (18S: NR_003286.4; 5.8S: NR_003285.2; 28S: NR_003287.4) was done by Bowtie2 (v2.3.5.1), with default parameters. Sorting, indexing and extraction of mapped reads was done using Samtools (v1.15.1), with option -F 4 to exclude the unmapped reads. Subsequent analysis was conducted using R: the quantification of ribosomal RNA (rRNA) 2′-O-Me residues was performed using the RNAmodR.RiboMethSeq package (v1.18.0), the final processed data were exported using the writexl (v1.5.0) package, display output were generated with the Prism software (v10.3.0).

### Pre-rRNA processing analysis by northern blotting

Total RNA was extracted from HEK293 cells labeled with or without 5eU with TRI reagent solution (AM9738, Thermofisher), according to the manufacturer’s instruction. 5 µg total RNA was resolved on a 1.2% agarose/6% formaldehyde denaturing gel, transferred overnight by capillarity onto a nylon Hybond N+ membrane (RPN203B, Cytiva), and hybridized with radioactively labeled probes. The probes were designed to detect all major pre-rRNA intermediates. The signal was acquired with a phosphorimager (FLA-7000, Fujifilm) and quantified with native multi gauge software (v3.1, Fujifilm).

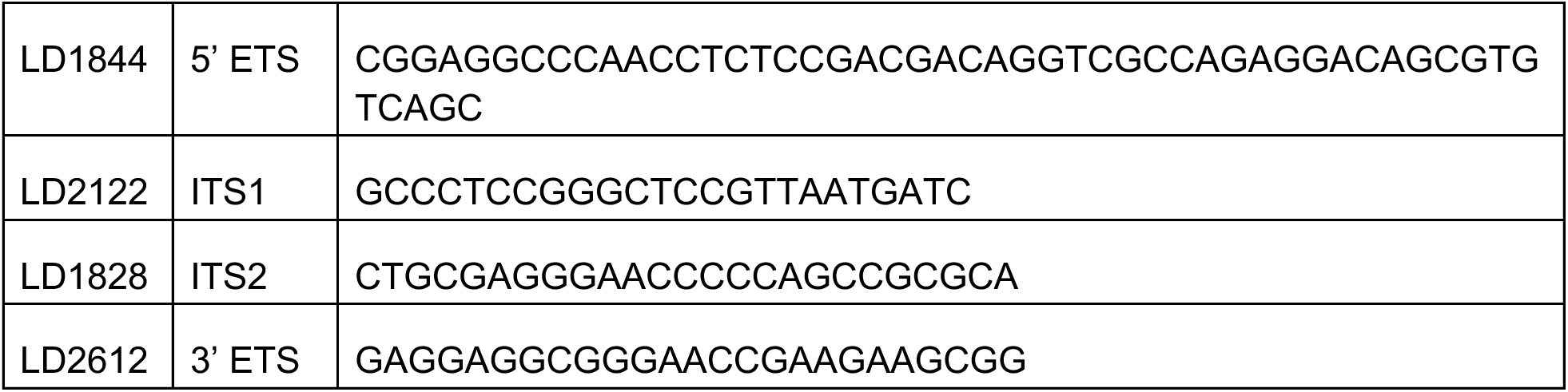

### Quantitative Image Analysis

All quantitative imaging measurements were performed with CellProfiler^74^ version 4.2.6. Cell segmentation utilized cellpose^75^ version 2.3.2 and the RunCellpose plugin for cellprofiler. Full computational methods, pipelines, and notebooks describing calculation of derived methods can be found at the github repository: https://github.com/SoftLivingMatter/image-analysis-quinodoz-jiang-2024. To deploy cellprofiler plugins on HPC systems, a snakemake workflow^76^ wrapper was developed which controls the resources and behavior of each invocation, which can be found at the github repository: https://github.com/softLivingMatter/snakemake-cellprofiler/.

#### Morphology

To quantitatively examine changes in cell morphology as a function of nucleolar perturbations, the following steps were performed in CellProfiler. First, the multichannel images were split into separate channels, such as GC, FC, DFC, FISH/5eU/other markers. The GC channel was used to define initial GC objects using minimum cross-entropy thresholding with a diameter range of 30-400 pixels, where each pixel corresponds to 0.0387 µm. Clumped objects were not separated as it produced too many falsely separated GC objects. Because the images do not have a nuclear or cytoplasmic stain and some metrics are best measured across an entire cell, we computationally assigned nucleoli to the same cell based if they were within 100 pixels to each other,producing a merged object mask. The mask was dilated by 50 pixels to measure metrics within the prospective nucleoplasm and find nucleolar features that may be outside the GC, such as FCs upon FVP treatment which causes detachment of the phases.

To find FC objects, the dilated GC objects were used to mask the FC image prior to performing “enhancing speckles” with a feature size of 20 pixels using the “Fast” setting in the EnhanceOrSuppressFeatures module. The FC objects were found in the enhanced FC image using an adaptive, three-class Otsu threshold with the middle intensity class assigned to the background. This was selected to ensure the identified FC objects were primarily in plane and focused. The adaptive window was 100 pixels, clumped objects were separated by intensity and shape, and FC objects were selected with diameters of 7-50 pixels. Since these FCs included objects within and outside of the GC, each FC was assigned to either a “nucleolar” or “extra-nucleolar” class using an overlap threshold of 50%, e.g. an FC which overlaps an initial GC object by more than 50% is considered a nucleolar FC.

DFC objects were defined by masking the DFC image with the dilated GC objects. Since the DFC phase organization is highly variable between treatments, the built-in “IdentifyPrimaryObjects” module wasn’t able to perform adequately in all cases. Instead, DFCs were found by first thresholding with an adaptive, three-class otsu cutoff with an 150 pixel window. The middle class was assigned to background to ignore the “diffuse” DFC phase that develops upon certain perturbations. The threshold image was converted to objects and then prospective DFCs were filtered to ensure they had an area of at least 20 pixels.

The GC, DFC, and FC objects were combined to a single mask which was used as the support for measuring Pearson’s correlation and overlap of the image channels. The size and shape of each object set was also measured and used for scaling some metrics, discussed below. The intensity of the FC channel was measured in FC objects, and GC and probe (e.g. FISH probe or 5eU) were measured in the GC and DFC phases. The distribution of FC and DFC intensity was measured over the initial GC objects using 20 scaled bins. A binary image of the initial GC objects was also measured to facilitate combining bins during post processing. Each object was related to its corresponding dilated GC object prior to exporting to a csv for further analysis in python.

The raw data produced by cellprofiler was further processed using jupyter notebooks and custom analysis scripts. Each object csv was read into a pandas dataframe and merged into a final result by the image and dilated GC unique identifier. Values such as total area or FC count were summed to provide a per-cell measurement. Average intensities per cell were calculated by first multiplying the mean intensity from cellprofiler with the object area, summing the result, then dividing by the total area per cell. Rim enrichment was calculated by summing the radial distribution fraction for the bins of interest and dividing by the fraction of the GC object over the same range of bins. Investigation of a range of rim widths found that the outer 20% of the GC rim provided the best discrimination between U3 knockdown and control cells. Circularity was estimated as 4*pi*area / perimeter^2.

#### Radial Distribution Function (RDF) Estimation

To facilitate broader utilization of RDF measurements, we developed a cellprofiler plugin “MeasureRDF” for calculating radial distributions of object sets, such as the intensities of FISH or 5eU signals within a 1 µm radius from the FC center, on a given input image. A separate plugin allows for further flexibility in performing upstream object finding and filtering and simplifies integration into other analysis tasks. In contrast to the “MeasureObjectIntensityDistribution” method available in cellprofiler, the “MeasureRDF” plugin utilizes a fixed distance in pixels and attempts to resolve overlapping objects as described below.

In the point-based measurement mode, the MeasureRDF plugin operates on a set of objects as the point sources along with a containing object. Here, we utilized the GC boundary as the masking objects and FCs as the point sources. For each GC, the set of FCs are considered together. The image intensities are mean centered with unit variance and pixel distances from each FC are determined. The RDF distribution is estimated by minimizing the difference between the scaled image intensity and a superposition of each FC point source with the same RDF. This allows for deconvolution of neighboring point sources while providing an averaged estimate of the RDF. The estimated intensities are rescaled to the original intensity units before reporting to the user. In all plots, the intensity is either “min-max” scaled between the minimum and maximum intensities in the RDF or “max” scaled by dividing by the maximum intensities in the RDF to highlight intensity dynamics.

The MeasureRDF plugin can also operate in a boundary-measurement mode, which was used for the SSU only analyses performed in Figure 4. In this setting, the source object boundaries are considered at r=0, defined as the radial position of 50% NOP56 normalized intensity, and the image intensity is measured as a function of distance from the boundary. The distance of each pixel to the object boundary is determined and used to estimate the RDF curve. In cases where neighboring point sources overlap, pixel intensities are assigned to the closest object.

#### Engineered plasmid measurements

For the images of *de novo* nucleoli, we manually classified individual nucleoli as “endogenous”, “de novo” or “hybrid” based on their intensities for plasmid-expressed 18S*/28S* RNA and endogenous 5’ ETS rRNA. Specifically, endogenous nucleoli were those with high endogenous 5’ ETS RNA FISH and no detectable plasmid-expressed 18S*/28S* RNA FISH signal. Conversely, de novo nucleoli were those with high 18S*/28S* plasmid RNA FISH signal, and no detectable endogenous 5’ ETS signal. Hybrid nucleoli contained FISH signals for both channels. Figure 3D demonstrates the range of nucleolar intensities of 5’ ETS and plasmid-expressed FISH intensities observed prior to further manual classification, where only high-confidence de novo vs. endogenous nucleoli were analyzed and any ambiguous nucleoli were excluded.

Since the SSU only rDNA plasmid produced nucleoli without detectable NPM1 intensity, a separate workflow was required to evaluate a subset of the morphology measurements described above. First, each channel was background corrected by subtracting the bottom 5 percentile value of each image. Next, the NOP56 channel was blurred with a 5-pixel sigma gaussian filter, which was found to produce better segmentation of SSU only nucleoli. The blurred NOP56 image was used to detect nucleoli with a two class, global Otsu threshold and diameters between 10 and 150 pixels without declumping. The initial nucleoli objects were dilated by 10 pixels to act as support for the rim-based RDF measurement. Each channel’s intensity was measured in the nucleoli and the 10-pixel rim as well as the object size and shape. For quality control, we manually checked the segmented SSU only nucleoli objects and excluded those that were out of focus or incorrectly segmented.

To measure cytoplasmic and nuclear intensities of the 18S* or 28S* RNA expressed from engineered plasmids, nuclei were segmented from the endogenous 28S FISH using cellpose with the nuclei model (inverted mask) and an expected object diameter of 300 pixels. The entire cell extents were then segmented from the endogenous 28S FISH stain (non-inverted mask) using the “cyto2” cellpose model and an object diameter of 500 pixels. The whole cell and nucleus objects were masked to ensure each cell had a nucleus and vice versa. Then the cell objects were masked with the nucleus to define the cytoplasm. The background subtracted endogenous 28S and engineered 18S* or 28S* RNA intensities were measured in the nuclei and cytoplasm.

## Supplementary Information

### Supplementary Videos (Attached)

Supplementary Video 1-2: Reformation of the multiphase nucleolus upon washout of flavopiridol (FVP). After 2 hours of 2 μM FVP treatment, cells were washed twice quickly with PBS and then imaged in regular medium. Images were acquired at 37 °C with 5% CO_2_, every two minutes after FVP removal over 90 min. Video 1 displays three layers of nucleolus (FC: RPA16-GFP; DFC: NOP56-mCherry; GC: mTagBFP2-NPM1) while Video 2 only displays FC and GC. Videos are registered through the HyperStackReg plugin from Fiji. Scale bars = 3 μm.

Supplementary Video 3: Video of images shown in Figure 5C. Modeling of SSU processing (5’ ETS cleavage) over time whereby an accumulation of SSU precursors (pre 5’ ETS cleavage) results in inversion of the nucleolar phases.

Supplementary Video 4: Video of images shown in Figure 5D. Modeling of RNA Pol I transcriptional inhibition. Decreased concentration of all SSU and LSU rRNA precursors results in the nucleolar capping morphology.

### Supplementary Notesa

Supplementary Note 1 (Attached): Kinetic modeling

Supplementary Note 2 (Attached): Nucleolus phase field modeling

Supplementary Note 3: We note that 18S* rRNA expressed from the SSU only plasmid exports successfully to the cytoplasm, suggesting that the GC may not be needed for SSU processing. However, we cannot exclude the possibility that these SSU particles undergo further processing in the GC of adjacent native nucleoli or hybrid nucleoli.

### Supplementary Table 1 (Attached)

RNA FISH probes, antibodies, RT-qPCR primers, ASOs/siRNAs and plasmids used in this study

