## Supplementary Note 1 for "Mapping and engineering RNA-controlled architecture of the multiphase nucleolus"

### Supplementary Note 1: kinetic model of transcription and processing of pre-ribosomal RNA

In this Supplementary Note, we present a chemical kinetics model to describe the transcription and processing of pre-ribosomal RNA (pre-rRNA) in the 5eU pulse-chase experiments. Fitting the model to the sequencing and imaging data allows us to extract effective reaction rates of transcription and processing, which can be used to infer steady-state properties such as the relative abundance of rRNA precursors.

#### Kinetic model for 5eU incorporation into pre-rRNA

In the 5eU pulse-chase experiments, the total amount of pre-rRNA can be measured by imaging the total 5eU intensity in the nucleus (Figure 1). We model the production of 5eU-labeled pre-rRNA with a two-step process: first, 5eU needs to be uptaken by the cell into the nucleus; second, the available 5eU is incorporated into pre-rRNA during transcription. Both steps are described by linear kinetics:

$$\frac{dc_0}{dt} = k_0 \Theta(-t)\Theta\left(t + t_{\text{pulse}}\right) - k_1 c_0$$

$$\frac{dc_1}{dt} = k_1 c_0$$

where $c_0$ is the amount of nuclear 5eU available for transcription, and $c_1$ is the amount of 5eU in pre-rRNA. $k_0$ and $k_1$ are the reaction rates of 5eU uptake and transcription, respectively. The Heaviside step functions $\Theta(t)$ and $\Theta\left(t + t_{\text{pulse}}\right)$ describe the pulse of 5eU, during $t \in \left(-t_{\text{pulse}}, 0\right)$.

The model can be solved analytically to obtain the total 5eU signal in the pre-rRNA:

$$c_1(t) = \begin{cases} 0 & t < -t_{\text{pulse}} \\ \frac{k_0}{k_1}\left(-1 + e^{-k_1(t+t_{\text{pulse}})} + k_1\left(t + t_{\text{pulse}}\right)\right) & -t_{\text{pulse}} \leq t < 0 \\ \frac{k_0}{k_1}\left(k_1 t_{\text{pulse}} - e^{-k_1 t} + e^{-k_1(t+t_{\text{pulse}})}\right) & t \geq 0 \end{cases}. \quad \text{[Eq.1]}$$

Eq.(1) agrees well with the total 5eU signal measured by imaging, with fitting parameters $k_0 = 0.052 \pm 0.006\ \text{min}^{-1}$ and $k_1 = 0.051 \pm 0.026\ \text{min}^{-1}$ (Figure 1). The fit allows us to extract the rate of transcription (of rRNA labeled by 5eU) $k_0 c_0$, with $c_0$ given by

$$c_0(t) = \begin{cases} 0 & t < -t_{\text{pulse}} \\ \frac{k_0}{k_1}\left(1 - e^{-k_1(t+t_{\text{pulse}})}\right) & -t_{\text{pulse}} \le t < 0 \\ \frac{k_0}{k_1} e^{-k_1 t}\left(1 - e^{-k_1 t_{\text{pulse}}}\right) & t \ge 0 \end{cases}. \quad \text{[Eq.2]}$$

#### Kinetic model for rRNA processing

Next, we consider the kinetics of rRNA processing as measured by sequencing in the pulse-chase experiments. For simplicity, we assume the processes to be limited by reaction rather than diffusion, which allows ignoring spatial degrees of freedom. We model each cleavage step as a first order reaction with rate $k_i$, with $i = 2,3,4,5$. $k_2$ corresponds to the cleavage of junctions 01 and 02, $k_3$ junctions 1 and 2, $k_4$ junction 3', and $k_5$ junction 4'. The abundance of individual pre-rRNA species is represented by $c_i$, with $i$ labeling the next cleavage step. The kinetic model is given by

$$\frac{dc_2}{dt} = k_1 c_0 - k_2 c_2, \quad \frac{dc_3}{dt} = k_2 c_2 - k_3 c_3,$$
$$\frac{dc_4}{dt} = k_3 c_3 - k_4 c_4, \quad \frac{dc_5}{dt} = k_4 c_4 - k_5 c_5, \quad \text{[Eq. 3]}$$
$$\frac{dc_6}{dt} = k_5 c_5,$$

where $c_1 = \sum_{i=2}^{5} c_i$ is the total amount of pre-rRNA. We fit the model to the cleavage fraction of each junction as measured by 5eU-sequencing:

$$f_{01} = \frac{c_3}{c_2 + c_3}, \quad f_{02} = 1 - \frac{c_2}{c_1}, \quad f_{1,2} = 1 - \frac{c_2 + c_3}{c_1},$$
$$f_{3'} = 1 - \frac{c_2 + c_3 + c_4}{c_1} = \frac{c_5 + c_6}{c_1}, \quad f_{4'} = \frac{c_6}{c_1},$$

where $f_i$ is the fraction of junction $i$ cleaved as measured by sequencing. Junctions 1 and 2 are averaged to obtain $f_{1/2}$. Junction 3' is not used in the fit. The definition of $f_{01}$ is slightly different from all the other fractions because the cleavage of junction 01 can no longer be detected once junction 1 is cut, while the cleavage of junction 2 is detectable in all the later species.

The kinetic model provides an excellent fit to the data (Figure 2). The best-fit parameters are $k_2 = 0.061\ \text{min}^{-1}$, $k_3 = 0.035\ \text{min}^{-1}$, $k_4 = 0.046\ \text{min}^{-1}$, and $k_5 = 0.046\ \text{min}^{-1}$. This also allows us to estimate the relative abundance of individual pre-rRNA species at steady state:

$$c_2^{ss} : c_3^{ss} : c_4^{ss} : c_5^{ss} \sim k_2^{-1} : k_3^{-1} : k_4^{-1} : k_5^{-1} = 19\% : 32\% : 25\% : 25\%. \quad \text{[Eq. 4]}$$

The abundance of 18S is slightly different since its processing does not involve cutting junctions 3' and 4'. We estimate that it is exported at rate $\tilde{k}_4 \sim 30$ min$^{-1}$ after the cleavage of junctions 1 and 2. The relative abundance of 18S is then given by

$$\tilde{c}_2^{ss}:\tilde{c}_3^{ss}:\tilde{c}_4^{ss} \sim k_2^{-1}:k_3^{-1}:\tilde{k}_4^{-1} = 22\%:38\%:40\%.$$

This provides an estimate of the fraction of processed versus unprocessed 18S in the multiphase reaction-diffusion model (Figure 5 in the main text).

#### Deconvolving the radial distribution of pre-rRNA

The kinetic model allows us to deconvolve the pre-rRNA distribution measured by 5eU imaging into the distribution of different pre-rRNA species. Let $\phi_i(r)$ be the steady-state radial distribution of pre-rRNA species $i$ (e.g. $i$ is a particular cleavage state), which is normalized by $\int_0^\infty \phi_i(r) 4\pi r^2 dr = 1$. Assuming that pre-rRNA processing is reaction limited (i.e. diffusion within each phase is much faster than cleavage), the radial distribution of the total pre-rRNA (since 5eU-imaging labels all the pre-rRNA species) $\phi(r,t)$ is given by a linear combination of the individual species:

$$\phi(r,t) = \frac{\sum_i c_i(t)\phi_i(r)}{\sum_i c_i(t)} = \sum_i \gamma_i(t)\phi_i(r),$$

where $\gamma_i(t) = \frac{c_i(t)}{\sum_i c_i(t)}$ is the fraction of pre-rRNA species $i$ at time $t$. $\phi(r,t)$ is normalized by $\int_0^\infty \phi(r,t) 4\pi r^2 dr = 1$ for all $t$.

Thus, the steady-state distribution of individual species can be obtained by solving a constrained non-negative least squares problem:

$$min_{\phi_i(r)} \sum_t \int_0^\infty \left(\phi(r,t) - \sum_i \gamma_i(t)\phi_i(r)\right)^2 4\pi r^2 dr, \quad \text{[Eq.5]}$$

subject to $\int_0^\infty \phi_i(r) 4\pi r^2 dr = 1$ and $\phi_i(r) \geq 0$ for $i$ running over all pre-rRNA species.

Here, we divide rRNA into early ($\phi_2$), middle ($\phi_3$), late ($\phi_4 + \phi_5$), and cytoplasmic ($\phi_6$) species. The relative abundance $\gamma_i(t)$ is predicted by the kinetic model (Figure 3A). $\phi(r,t)$ is measured by 5eU imaging (Figure 3B). Solving the optimization problem gives the radial distribution function $\phi_i(r)$ (Figure 3C) and probability distribution function $4\pi r^2 \phi_i(r)$ (Figure 3D), which demonstrate that pre-rRNA processing correlates with its outward movement.

A similar deconvolution can be done to the RNA FISH measurements. The normalized FISH signal for junction $i$ is given by

$$y_i(r) = \frac{\sum_{\{j \le i\}} c_j^{ss} \phi_j(r)}{\sum_{\{j \le i\}} c_j^{ss}}, \text{ [Eq. 6]}$$

where $c_j^{ss}$ is the steady-state abundance of species $j$ given in Eq. (4). Solving Eq. (6) for $\phi_i(r)$ gives the spatial distribution of intermediates predicted by FISH, which can be compared with that from 5eU-seq and imaging (Figure 4).

#### Figures

Figure 1: Fitting the total 5eU signal in the nucleus to the model (Eq. (1)). The fit only used data in the first two hours after the pulse.

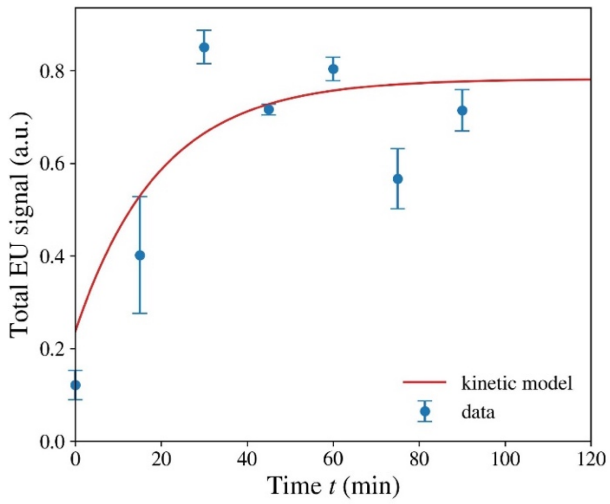


Figure 2: The kinetic model (Eq. (3)) captures the cleavage fraction of nascent rRNA as measured by 5eU-sequencing.

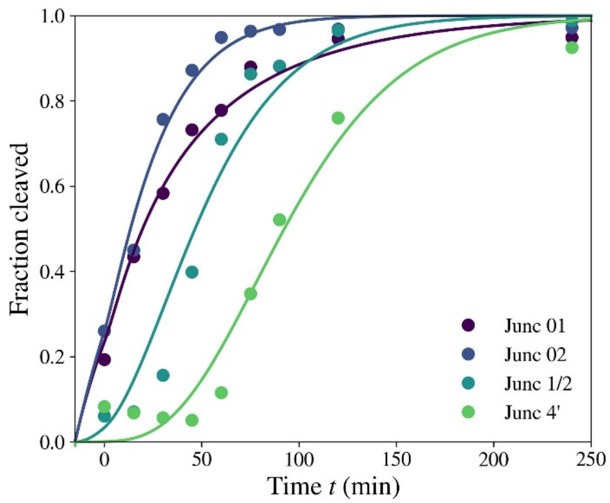

Figure 3: Deconvolving the radial distribution of nascent rRNA

(A) rRNA abundance $\gamma_i(t)$ as a function of time, obtained from the kinetic model.

(B) the radial distribution of nascent rRNA at different times, obtained by normalizing the 5eU imaging data.

(C—D) the radial distribution function $\phi_i(r)$ (C) and probability distribution function $4\pi r^2\phi_i(r)$ (D) of individual rRNA species, inferred by solving the optimization problem Eq. (5).

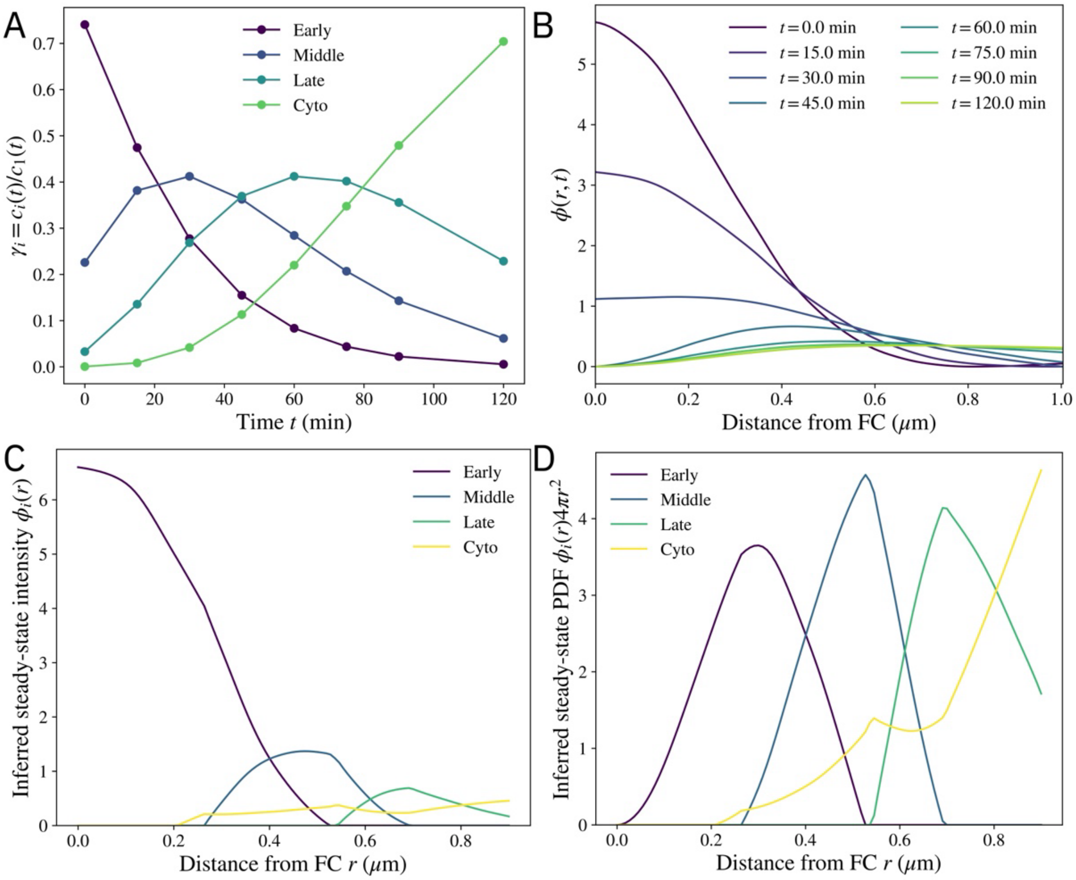

Figure 4: The radial distribution of early, middle, and late pre-rRNA species as inferred from 5eU-imaging (top) and FISH (bottom). For 5eU-imaging, the distribution is inferred by solving the optimization problem Eq. (5). For FISH, the spatial distribution is obtained by solving Eq. (6) for $\phi_i(r)$. The radius of the circles is 0.9 $\mu m$.

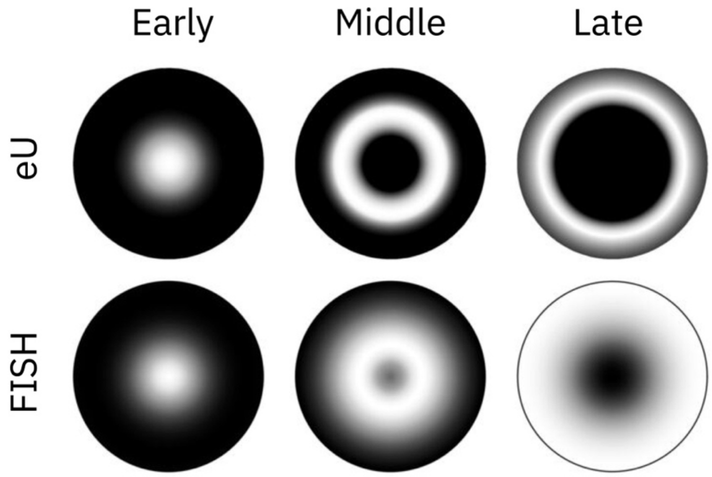
