## Supplementary Note 2 for "Mapping and engineering RNA-controlled architecture of the multiphase nucleolus"

### A. Thermodynamic model

Given the experimental evidence of nucleolar morphology changes upon perturbations to rRNA composition, we seek to develop a model that captures the change in the surface tensions between nucleolar phases as well as the changes in the partitioning of rRNA in the different phases upon rRNA perturbations. We first consider the Flory-Huggins model, a well-established thermodynamic model that describes polymer phase separation. To capture the essential physics, the system consists of the nucleolar species [DFC, GC, and nucleoplasm (NP)], and the rRNA species [SSU before 5'ETS cleavage, i.e. unprocessed 18S (u18S), SSU after 5'ETS cleavage, i.e. processed 18S (p18S), and 28S]. We denote the set of nucleolar species and nucleoplasm as $N = \{DFC, GC, NP\}$ and the set of rRNA species as $R = \{u18S,\ p18S,\ 28S\}$. The free energy density of the mixture g follows

$$\frac{g}{k_B T} = \sum_i \frac{1}{N_i} \phi_i \ln \phi_i + \frac{1}{2} \sum_{ij} \chi_{ij} \phi_i \phi_j, \qquad (1)$$

where $k_B$ is the Boltzmann constant, T is the temperature; $N_i$ and $\phi_i$ are the degree of polymerization and the volume fraction of component i, respectively. The volume fractions satisfy the normalization condition $\sum_i \phi_i = 1$. The summation indices i and j iterate over all chemical components mentioned above. For simplicity, we set all degrees of polymerization to be equal to $N_0$. In the free energy density in the above Eq. (1), the first and second terms correspond to the entropy and enthalpy of mixing, respectively. $\chi_{ij}$ is the interaction parameter between components i and j – a higher value means stronger repulsion between the two, and $\chi_{ij} = \chi_{ji}\ (i \neq j)$, $\chi_{ii} = 0$. This determines the phase separation between nucleolar compartments as well as the affinity of rRNA to each nucleolar phase. For example, if $\chi_{GC,28S} < \chi_{NP,28S}$, then 28S is more enriched in the GC compared to the nucleoplasm. When $\chi_{ij}$ is sufficiently high, the system undergoes phase separation into a phase that is rich in component i and another coexisting phase that is rich in component j. Following the Cahn-Hilliard theory of phase separation, the free energy of such a spatially heterogeneous system where phase separation occurs is

$$G = c_0 \int \left( g + \frac{1}{2} \kappa \sum_i |\nabla \phi_i|^2 \right) dV, \qquad (2)$$

where the gradient term corresponds to the energy of interaction at the interface, and $c_0$ is the number concentration of monomers. The phase equilibrium condition corresponds to the minimization of the free energy subject to the mass conservation of all chemical components. This is equivalent to a uniform chemical potential $\mu_i$, which is defined to be the variational derivative of the free energy with respect to the volume fraction,

$$\mu_i \equiv \frac{1}{c_0} \frac{\delta G}{\delta \phi_i} = \mu_i^0. \qquad (3)$$

Suppose that phases α and β form a flat interface in an infinitely large domain, the surface tension between the two phases $\gamma_{\alpha\beta}$ is defined to be the excess energy associated with the interface joining the two bulk phases,

$$\gamma_{\alpha\beta} = c_0 \int_{-\infty}^{\infty} \left( g + \frac{1}{2}\kappa \sum_i |\nabla\phi_i|^2 - \sum_i \mu_i^0 (\phi_i - \phi_i^{(\alpha)}) \right) dx, \qquad (4)$$

where x is in the direction normal to the interface, $\phi_i$ is the volume fraction at equilibrium, $\phi_i^{(\alpha)}$ is the volume fraction in phase α away from the interface.

It is known that when $\chi_{ij}$ is large, the surface tension between the two phases is approximately proportional to $\chi_{ij}$ (Mao et al. 2019, Mao et al. 2020). However, it is unclear how the presence and partitioning of rRNA species affect the surface tension. Therefore, we numerically calculate the surface tension by performing a phase field simulation that finds the equilibrium state. Specifically, we perform one-dimensional simulations with periodic boundary conditions and find an equilibrium state where three phases, which are rich in DFC, GC, and NP, separately, coexist. We then calculate the integral in Eq. (4) between all bulk phases to find the surface tensions.

In the following text, we use $\gamma_{DFC,GC}$ to denote the surface tension between the DFC and GC phases, and similarly $\gamma_{DFC,NP}$, $\gamma_{GC,NP}$. Under normal conditions, the DFC phase is surrounded by the GC phase, indicating that the surface tensions satisfy the inequality $\gamma^{(1)}_{DFC,NP} \geq \gamma^{(1)}_{DFC,GC} + \gamma^{(1)}_{GC,NP}$, where the superscript (1) indicates a system (1) where the concentration of processed 18S is high, unprocessed 18S is low, and that of the 28S is normal. Under U3 ASO treatment and treatment with mutant plasmids, the GC phase is surrounded by the DFC phase, indicating that $\gamma^{(2)}_{GC,NP} \geq \gamma^{(2)}_{DFC,GC} + \gamma^{(2)}_{DFC,NP}$, where the superscript (2) indicates a system (2) where the concentration of processed 18S is low, unprocessed 18S is high, and that of the 28S is normal, and the average volume fractions of DFC and GC are equal to that of the system (1).

Therefore, in order to determine if a set of parameters $\chi_{ij}$ exists that can recapitulate these two morphologies, we perform the following optimization

$$\min_{\chi_{ij}} \Gamma(\gamma^{(1)}_{DFC,GC} + \gamma^{(1)}_{GC,NP} - \gamma^{(1)}_{DFC,NP}) + (1 - \Gamma)(\gamma^{(2)}_{DFC,GC} + \gamma^{(2)}_{DFC,NP} - \gamma^{(2)}_{GC,NP}), \qquad (5)$$

where $0 < \Gamma < 1$ is a weighting factor. In the optimization, the interaction parameters between rRNA species is set to 0, i.e., $\chi_{i\in R, j\in R} = 0$, while the interaction between nucleolar components $\chi_{i\in N, j\in R}$ and between nucleolar and rRNA species $\gamma_{i\in N, j\in R}$ are free variables. The optimization results show that while it is possible to achieve full wetting in one system and partial wetting in the other (e.g., $\gamma^{(1)}_{DFC,GC} + \gamma^{(1)}_{GC,NP} = \gamma^{(1)}_{DFC,NP}$ and $\gamma^{(2)}_{DFC,GC} + \gamma^{(2)}_{DFC,NP} < \gamma^{(2)}_{GC,NP}$), it fails to find a set of parameters that achieve full wetting in both systems with different orderings of DFC and GC. This indicates that the pairwise Flory-Huggins model is insufficient to explain the change of nucleolus from a normal to an inverted morphology as a result of rRNA processing.

These results motivate us to consider a higher-order model that includes the interaction terms between three components. This correction is not merely a mathematical construction but also stems from the physics of three-body interaction that is reflected in the third virial coefficient. Specifically, we consider the following free energy density

$$\frac{g}{k_B T} = \sum_{i \in N \cup R} \frac{1}{N_i} \phi_i \ln \phi_i + \frac{1}{2} \sum_{ij \in N, k \in R} (\chi_{ij} + w_{ijk} \phi_k) \phi_i \phi_j + \sum_{i \in N, k \in R} \chi_{ik} \phi_i \phi_k. \quad (6)$$

The new three-body term indicates that rRNA species can directly control the effective interaction between two nucleolar compartments and hence the surface tension. When $w_{ijk} > 0$ ($w_{ijk} < 0$), rRNA species k increases (decreases) the surface tension between i-rich and j-rich phases. The pairwise interaction between rRNA and nucleolar species dictates the partitioning of rRNA. For simplicity, we set the degrees of polymerization as $N_i = N_n,\ i \in N$ and $N_i = N_r,\ i \in R$. To simplify the model, we assume that the concentration of rRNA is low compared to that of the nucleolar components and define normalized rRNA volume fraction to be $\tilde{\phi}_k = \phi_k / \phi_0$, where $\phi_0$ is a characteristic volume fraction of rRNA such that $\tilde{\phi}_k \sim O(1)$ and $\phi_0 \ll 1$. As a result, the normalization condition for the volume fractions is approximately $\sum_{i \in N} \phi_i = 1 - \phi_0 \sum_{k \in R} \tilde{\phi}_k \sim O(1)$. We then define normalized interaction parameters $\tilde{\chi}_{ij} = \chi_{ij} N_n\ (i, j \in N)$, $\tilde{w}_{ijk} = w_{ijk} \phi_0 N_n\ (i, j \in N, k \in R)$, $\tilde{\chi}_{ik} = \chi_{ik} N_r\ (i \in N, k \in R)$. Therefore, Eq. (6) becomes

$$\frac{g}{k_B T} = \frac{1}{N_n} \left( \sum_{i \in N} \phi_i \ln \phi_i + \frac{1}{2} \sum_{ij \in N, k \in R} (\tilde{\chi}_{ij} + \tilde{w}_{ijk} \tilde{\phi}_k) \phi_i \phi_j \right) + \frac{\phi_0}{N_r} \left( \sum_{k \in R} \tilde{\phi}_k \ln(\phi_0 \tilde{\phi}_k) + \sum_{i \in N, k \in R} \tilde{\chi}_{ik} \phi_i \tilde{\phi}_k \right). \quad (7)$$

Note that the term involving $\ln \phi_0$ does not influence the phase diagram and surface tension because it is a linear term with respect to $\tilde{\phi}_k$. For simplicity, we set $\frac{1}{N} = \frac{\phi_0}{M}$.

Because $\tilde{\chi}_{ij}\ (i, j \in N)$ and $\tilde{w}_{ijk}\ (i, j \in N, k \in R)$ primarily control the surface tensions and $\tilde{\chi}_{ik}\ (i \in N, k \in R)$ primarily controls the affinity of rRNA to nucleolar components, these interaction parameters can be set based on the experimentally observed nucleolus morphology and rRNA partitioning. Table 1 displays the parameters used in our subsequent simulations.

| Pairwise interaction parameters between nucleolar species $\tilde{\chi}_{ij}\ (i, j \in N)$ | | | |
|---|---|---|---|
| (i, j) | (DFC, NP) | (GC, NP) | (DFC, GC) |
| | 3 | 2.1 | 2.5 |
| Three-body interaction parameters between rRNA and nucleolar species $\tilde{w}_{ijk}\ (i, j \in N, k \in R)$ | | | |

| k \ (i,j) | (DFC, NP) | (GC, NP) | (DFC, GC) |
| --- | --- | --- | --- |
| Unprocessed 18S | -0.6 | 1 | -1.6 |
| Processed 18S | 2.3 | -2.5 | -0.6 |
| 28S | 0 | 1 | 1 |
| Pairwise interaction parameters between rRNA and nucleolar species $\tilde{\chi}_{ik}$ $(i \in N, k \in R)$ | | | |
| | DFC | GC | NP |
| Unprocessed 18S | -1 | -0.4 | 0 |
| Processed 18S | 0.8 | -1 | 0 |
| 28S | 0 | -1.05 | 0.15 |

Table 1 Interaction parameters used in the simulations

Because rRNA is soluble and has low concentrations, we omit the contribution of the energy associated with its gradient. Hence for a spatially heterogeneous system, the total free energy is

$$G = c_0 \int \left( g + \frac{1}{2} \kappa \sum_{i \in N} |\nabla \phi_i|^2 \right) dV \qquad (8)$$

We define normalized quantities, $\tilde{g} = gN/(c_0 k_B T)$, $\tilde{G} = GN/(c_0 k_B T)$, $\tilde{\kappa} = \kappa N/(k_B T)$, and the dimensionless chemical potentials

$$\tilde{\mu}_i \equiv \frac{\delta \tilde{G}}{\delta \phi_i} = \frac{\partial \tilde{g}}{\partial \phi_i} - \frac{\partial \tilde{g}}{\partial \phi_{NP}} - \tilde{\kappa} \left( \nabla^2 \phi_i - \nabla^2 \phi_{NP} \right) (i = DFC, GC), \quad (9)$$

$$\tilde{\mu}_k \equiv \frac{\delta \tilde{G}}{\delta \tilde{\phi}_k} = \frac{\partial \tilde{g}}{\partial \tilde{\phi}_k} \ (k \in R), \qquad (10)$$

where the Laplacian term involving $\phi_{NP}$ is due to the normalization condition $\phi_{NP} = 1 - \phi_{DFC} - \phi_{GC}$.

Based on the parameters in Table 1, we calculate the surface tension between phases at equilibrium for some typical average compositions that represent the normal, inversion, transcription-inhibited, and SSU-only plasmid phenotypes, as shown in Table 2. Note that in the normal phenotype (wild type), the ratio of unprocessed 18S to processed 18S rRNA is 0.1:0.9, consistent with the kinetic modeling results. The average volume fractions of the nucleolar components are kept the same across all simulations. From Table 2, we see that the normal phenotype satisfies $\gamma_{DFC,NP} = \gamma_{DFC,GC} + \gamma_{GC,NP}$. In 1D simulations, this corresponds to a complete GC wetting at the DFC-NP interface. In contrast, the inversion phenotype satisfies $\gamma_{GC,NP} = \gamma_{DFC,GC} + \gamma_{DFC,NP}$, which corresponds to a complete DFC wetting at the GC-NP interface. Moreover, we find that $\gamma_{DFC,NP}$ and $\gamma_{GC,NP}$ is higher for the inversion phenotype than the normal phenotype, consistent with the observation that the nucleoli fuse upon U3 ASO treatment. It is also consistent with the experimental observation that the nucleoli under the mutant SSU-only plasmid where 5'ETS cleavage is disrupted have a higher sphericity than the normal SSU-only plasmid. In contrast, $\gamma_{DFC,GC}$ is lower for the inversion phenotype than the normal phenotype, and the DFC phase in the inversion phenotype shows more DFC-GC mixing than that in the normal phenotype, consistent with experimental observations. The SSU-only plasmid does not have a GC phase, instead, GC is about equally distributed in the DFC and NP phases.

| Typical average composition | Normal phenotype | Inversion phenotype | Transcription inhibited | SSU only |
|---|---|---|---|---|
| $\phi_{DFC}$ | 0.12 | 0.12 | 0.12 | 0.12 |
| $\phi_{GC}$ | 0.32 | 0.32 | 0.32 | 0.32 |
| $\tilde{\phi}_{u18S}$ | 1.5 | 1 | 0.18 | 0.6 |
| $\tilde{\phi}_{p18S}$ | 1 | 0.01 | 0.12 | 0.4 |
| $\tilde{\phi}_{28S}$ | 1 | 0.75 | 0.3 | 0.01 |
| Normalized surface tension | Normal morphology | Inversion morphology | Transcription inhibited | SSU only |
| $\gamma_{DFC,NP}$ | 0.755 | 0.769 | 0.240 | 0.311 |
| $\gamma_{GC,NP}$ | 0.302 | 0.970 | 0.194 | NaN |
| $\gamma_{DFC,GC}$ | 0.453 | 0.201 | 0.119 | NaN |
| Equilibrium composition ($\phi_{DFC}$, $\phi_{GC}$) | Normal morphology | Inversion morphology | Transcription inhibited | SSU only |
| DFC phase | 0.589, 0.406 | 0.516, 0.464 | 0.738, 0.170 | 0.577, 0.383 |
| GC phase | 0.002, 0.660 | 0.028, 0.966 | 0.089, 0.742 | NaN |
| NP phase | 0.006, 0.032 | 0.032, 0.030 | 0.075, 0.178 | 0.023, 0.307 |
| Equilibrium composition ($\tilde{\phi}_{u18S}$, $\tilde{\phi}_{p18S}$, $\tilde{\phi}_{28S}$) | Normal morphology | Inversion morphology | Transcription inhibited | SSU only |

| | | | | |
|---|---|---|---|---|
| DFC phase | 3.262, 0.579, 0.901 | 1.959, 0.009, 0.628 | 0.396, 0.045, 0.230 | 1.352, 0.212, 0.010 |
| GC phase | 1.096, 1.806, 1.304 | 1.040, 0.019, 1.420 | 0.205, 0.193, 0.440 | NaN |
| NP phase | 1.040, 0.587, 0.819 | 0.687, 0.007, 0.533 | 0.151, 0.100, 0.255 | 0.441, 0.440, 0.010 |

Table 2. Equilibrium properties of different phenotypes. The normalized surface tension is defined to be $\tilde{\gamma}_{ij} = \gamma_{ij}/\left(c_0 k_B T N^{-1}\sqrt{\tilde{\kappa}}\right)$

### B. Dynamic model

Next, we proceed to model the nonequilibrium process in the nucleolus. To capture the essential physics, we consider and simplify two major pathways of rRNA synthesis that produce SSU and LSU in parallel. The SSU pathway is simplified to consist of one intermediate, that is, an unprocessed 18S is produced and then converted into a processed 18S in the DFC with rate coefficients $k_{0,18S}$ and $k_c$, respectively. The LSU pathway omits all processing steps, i.e. 28S is produced in the DFC with a rate coefficient $k_{0,28S}$. The production rates for the SSU and LSU pathways are assumed to be equal ($k_{0,18S} = k_{0,28S} = k_0$) except for the SSU-only plasmid, for which there is no 28S production. All rRNA species degrade and are exported from the NP with rate constant $k_{d,i}$. All reactions are assumed to follow first-order reaction kinetics. All components are assumed to have the same normalized mobility constant L. The normalization of the mobility constant absorbs all constants from normalizing the chemical potential. DFC and GC nucleolar species are chemically inert and they follow

$$\frac{\partial \phi_i}{\partial t} = L\nabla^2 \tilde{\mu}_i \ (i = DFC,\ GC). \quad (8)$$

The normalized volume fraction of rRNA components satisfy

$$\frac{\partial \tilde{\phi}_{u18S}}{\partial t} = L\nabla^2 \tilde{\mu}_{u18S} + k_{0,18S}\phi_{DFC} - k_c \phi_{DFC}\tilde{\phi}_{u18S} - k_{d,u18S}\phi_{GC}\tilde{\phi}_{u18S}, \quad (9)$$

$$\frac{\partial \tilde{\phi}_{p18S}}{\partial t} = L\nabla^2 \tilde{\mu}_{p18S} + k_c \phi_{DFC}\tilde{\phi}_{u18S} - k_{d,p18S}\phi_{GC}\tilde{\phi}_{p18S}, \quad (10)$$

$$\frac{\partial \tilde{\phi}_{28S}}{\partial t} = L\nabla^2 \tilde{\mu}_{28S} + k_{0,28S}\phi_{DFC} - k_{d,28S}\phi_{GC}\tilde{\phi}_{28S}. \quad (11)$$

We define a reaction-diffusion length scale based on the production of 18S and 28S, $l_d = \sqrt{L/(k_0 \phi^{(DFC)}_{DFC})}$, where $\phi^{(DFC)}_{DFC}$ is the volume fraction of DFC nucleolar species in the DFC phase in the normal phenotype as tabulated in Table 2. Given the reaction-limited assumption in the kinetic modeling, we set $l_d/L_0 = 5$, where $L_0$ is the size of the domain. We define the characteristic time scale $t_* = (k_{d,28S}\phi^{(GC)}_{GC})^{-1}$, where $\phi^{(GC)}_{GC}$ is the volume fraction of

GC nucleolar species in the GC phase in the normal phenotype as tabulated in Table 2. The characteristic interfacial width relative to the domain size is set to be $\sqrt{\kappa}/L_0 = 0.02$. The relative ratios of the rate constants in the normal phenotypes are determined by the overall mass balance based on the equilibrium states computed based on the average composition in Table 2. The transcription-inhibited phenotype has a lower production rate than the normal phenotype. The 28S degradation rate is lowered slightly to prevent dissolution of the GC phase due to the lack of 28S. Compared to the normal phenotype, all rate constants of the inversion phenotype remain the same except for the processing rate constant $k_c = 0$, all rate constants of the SSU-only plasmid remain the same except for the production rate of 28S $k_{0,28S} = 0$.

| Phenotype | 18S production rate constant $k_{0,18S}/k_0$ | 28S production rate constant $k_{0,28S}/k_0$ | 18S processing rate constant $k_c/k_0$ |
|---|---|---|---|
| Normal | 1 | 1 | 0.024 |
| Inversion | 1 | 1 | 0 |
| Transcription inhibited | 0.235 | 0.235 | 0.024 |
| SSU only | 1 | 0 | 0.024 |
| Phenotype | Unprocessed 18S degradation rate constant $k_{d,u18S}/k_0$ | Processed 18S degradation rate constant $k_{d,p18S}/k_0$ | 28S degradation rate constant $k_{d,28S}/k_0$ |
| Normal | 0.295 | 0.019 | 0.234 |
| Inversion | 0.295 | 0.019 | 0.234 |
| Transcription inhibited | 0.295 | 0.019 | 0.181 |
| SSU only | 0.295 | 0.019 | 0.234 |

Table 3 Reaction rate constants used in the following dynamics simulations

Based on the parameters above, we perform simulations for the different phenotypes based on the average composition in Table 2 and obtain the steady-state solution as shown in Fig. 1. The results agree with the morphology and rRNA partitioning observed in experiments.

Next, we perform perturbation simulations that cause the system to transition from one state to the other. In one case, we start from the steady state of the normal phenotype as the initial condition and at t = 0, the rate constants are switched to those of the inversion phenotype, in order to simulate the process of U3 ASO treatment. Fig. 2 shows that because the 18S processing is inhibited, the average concentration of unprocessed 18S increases while that of the processed 18S decreases, correspondingly the snapshots below show that the DFC phase first moves to the edge of the GC, and then gradually envelops GC. In another case, we start from the steady state of the normal phenotype and switch the rate constants to those of the transcription-inhibited phenotype to simulate the CX treatment. Fig. 3 shows that the concentrations of all rRNA species decrease and the simulation shows that DFC moves to the edge of the GC.

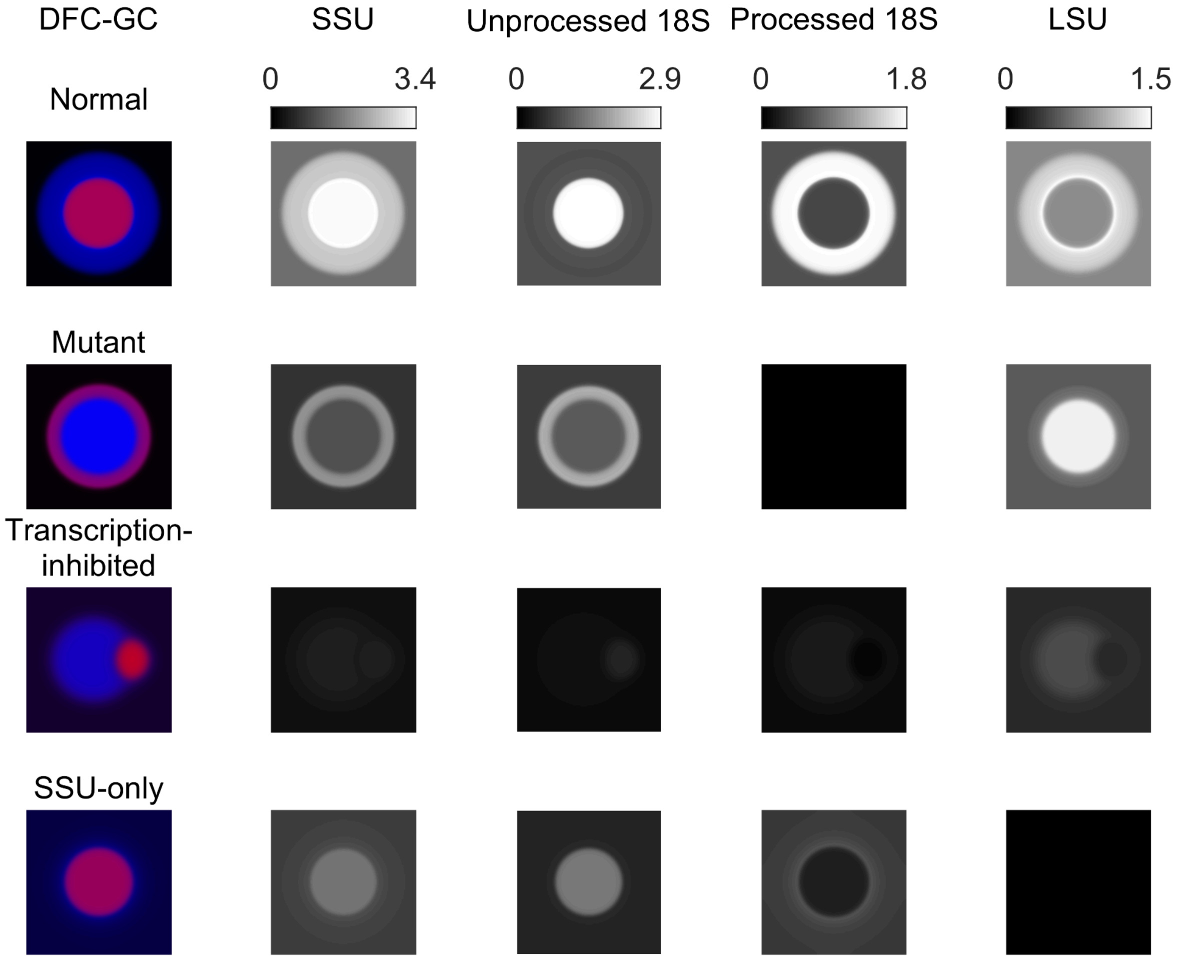


Figure 1. Nucleolus simulation (based on Eqs. (9)-(11)) at nonequilibrium steady state for some typical phenotypes whose average compositions are given in Table 2. From left to right, the images are DFC-GC merged (where the RGB values are $R = \phi_{DFC}$, $G = 0$, $B = \phi_{GC}$), total SSU normalized concentration ($\widetilde{\phi}_{u18S} + \widetilde{\phi}_{p18S}$), the normalized concentrations of unprocessed and processed 18S, and that of 28S (LSU), separately.

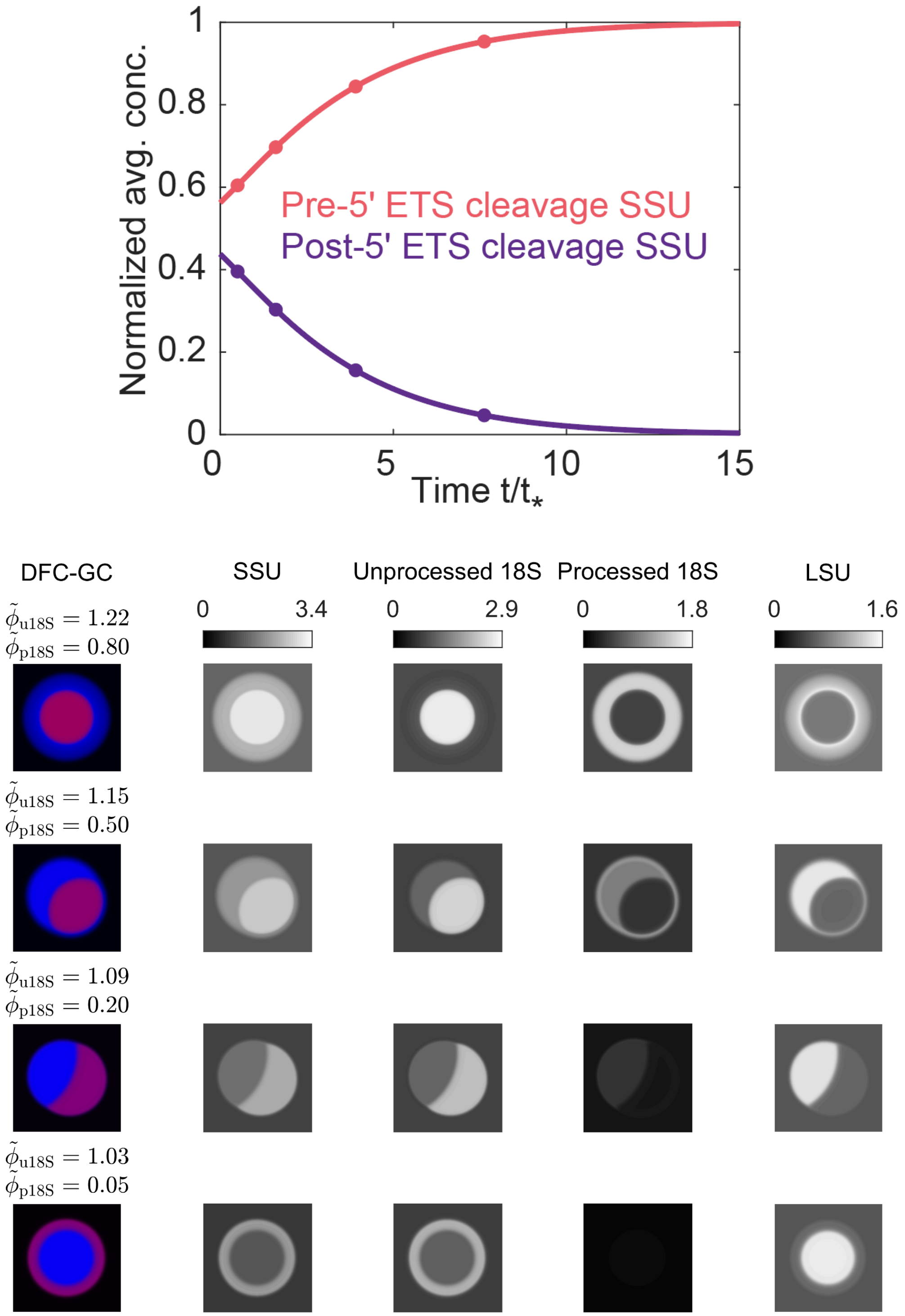


Figure 2 (a) Evolution of the normalized average concentration of unprocessed and processed 18S over time. The system is initialized at the normal phenotype steady state as shown in Fig. 1. At t = 0, the processing 18S is turned off. (b) Snapshots of the nucleolus morphology and rRNA distribution over time from top to bottom. Snapshots are taken at time points that correspond to the circles in (a).

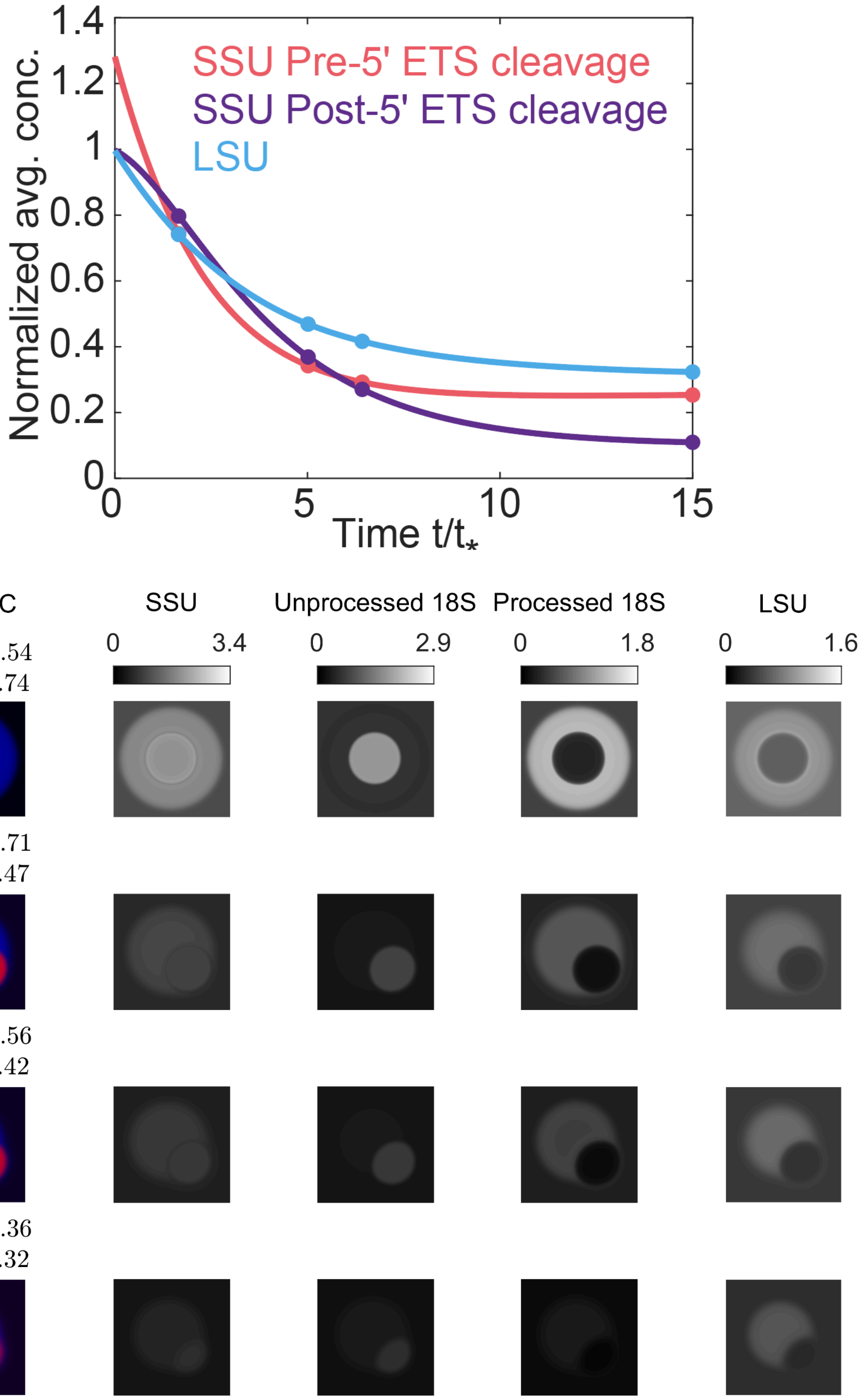


Figure 3 (a) Evolution of the normalized average concentration of SSU ($\tilde{\phi}_{u18S} + \tilde{\phi}_{p18S}$) and 28S over time. The system is initialized at the normal phenotype steady state as shown in Fig. 1. At t = 0, the production of 18S and 28S is turned off. (b) Snapshots of the nucleolus morphology and rRNA distribution over time from top to bottom. Snapshots are taken at time points that correspond to the circles in (a)
